# BioScientist Agent: Designing LLM-Biomedical Agents with KG-Augmented RL Reasoning Modules for Drug Repurposing and Mechanistic of Action Elucidation

**DOI:** 10.1101/2025.08.08.669291

**Authors:** Fan Zhang, Yalong Zhao, Weihan Zhang, Lipeng Lai

## Abstract

Drug discovery is protracted, resource-intensive, and afflicted by attrition rates exceeding 90 %, which leaves most diseases, particularly rare or neglected indications, without effective therapies. Drug repurposing offers a cost effective alternative, yet systematic identification of novel drug indication pairs and mechanistic rationales remains hindered by the scale and heterogeneity of biomedical knowledge. We present **BioScientist Agent**, an end to end framework that unifies a billion-fact biomedical knowledge graph with (i) a variational graph auto-encoder for representation learning and link prediction driven drug repurposing, (ii) a reinforcement learning module that traverses the graph to recover biologically plausible mechanistic paths, and (iii) A large language model (LLM) multi-agent layer that orchestrates these components, enabling inference of target pathways for a drug disease pair, and automatic generation of coherent causal reports. In all downstream tasks, the BioScientist Agent surpasses existing state of the art baseline models across various metrics and provides mechanistic explanations consistent with the literature. Its open and modular design accelerates hypothesis generation and reduces experimental overhead in early stage discovery.

## 1 Introduction

Drug discovery is a time-consuming and capital-intensive endeavor, typically taking over ten years and costing more than $2 billion per approved entity, yet more than 90% of clinical candidates fail. Consequently, thousands of common, rare, and neglected diseases remain without effective treatment. The clinical efficacy of drug therapies is often limited by resistance and severe toxic side effects, creating an urgent need for new treatments. A promising strategy that has gained attention in recent years is drug repurposing. This involves systematically identifying new indications for molecules with established safety profiles. Repurposed drugs are often cost-effective and proven safe, significantly accelerating the drug development process [XSHJ24]. Successful candidates include chlorambucil and bufulfone, which were originally developed as alkylating agents based on the toxic chemical warfare agent mustard gas but were later found effective for treating leukemias [Sch21]. Similarly, thalidomide, despite causing severe birth defects, has been repurposed to treat leprosy and multiple myeloma [BIS^+^22]. Additionally, arsenic trioxide and all-trans retinoic acid, a metabolite of vitamin A, are examples of other compounds approved by the FDA in 2000 for the treatment of acute promyelocytic leukemia [RLZ13]. Despite the high research and application value of drug repurposing, few drugs have successfully transitioned to clinical application through this method. It requires approaches capable of mining the vast and heterogeneous biomedical knowledge space and providing transparent mechanistic evidence.

Knowledge graphs (KGs) are crucial tools for drug repurposing research because they encompass comprehensive information on drugs, diseases, and related cellular experiments, genes, and protein targets. They can assess the potential of different drugs to treat diseases and construct the Mechanism of Action (MoA) through the connections between various types of nodes. By evaluating the reliability of the MoA with existing experimental research, KGs can enhance the accuracy of drug repurposing and improve the efficiency of drug applications. Knowledge graphs can extract, organize, and effectively manage knowledge from large-scale data to enhance the quality of information services and provide users with more intelligent services. Currently, research has developed extensive knowledge graphs for biological data, such as BioKG [WMN20], CKG [SCN^+^22], and RTX-KG2 [WGK^+^22], for researchers to perform data mining and analysis. These graphs integrate nearly all biomedical databases, such as DrugBank [KWK^+^24], ChEMBL [GBB^+^12], Gene Ontology (GO) [ABB^+^00], UniProt [Con19], and the Human Metabolome Database (HMDB) [WGO^+^22]. This extensive integration greatly benefits various biomedical studies, including drug repurposing [GDX22, MZLK23], adverse drug reaction analysis [ZSD^+^21, JMM22], and proteomics data analysis [SCN^+^22].

Simultaneously, large language models (LLMs) have evolved into versatile reasoning engines capable of planning, tool use, and scientific discourse. Combined with structured knowledge graph (KG) representations, LLMs can encode the semantic content of node descriptions into vector representations, facilitating tasks such as semantic textual similarity, information retrieval, and clustering [BAM^+^24]. Furthermore, Retrieval-Augmented Generation (RAG) technology enhances the reasoning capabilities of large models. When a query is received, the system first searches for the most relevant documents in an external corpus and then generates responses based on these retrieved facts. This design allows for more credible and verifiable outputs. In the biomedical field, RAG enables models to provide citations for any factual statements they make, such as references to medical textbooks, clinical guidelines from authoritative bodies, or peer reviewed scientific research. However, failures in retrieval quality or integration can still introduce inaccuracies, highlighting the need for robust validation even in a retrieval-augmented environment, as issues of hallucinations and faulty reasoning persist [SZZ^+^24, HC22]. We aim to leverage information extraction from biological knowledge graphs and combine symbolic reasoning with natural language synthesis using LLM-based agents. This approach will automate the generation of drug mechanisms of action and enhance interpretability.

Against this backdrop we introduce **BioScientist Agent**, a KG-augmented, LLM-orchestrated platform for drug repurposing and mechanistic elucidation. Our principal contributions are as follows:

1. We curate and preprocess RTX-KG2 (v2.9.2) to obtain a 6.3 million-node, 41 million-edge biomedical KG and train a variational graph auto-encoder (VGAE) that achieves state-of-the-art performance on drug–disease link-prediction benchmarks.
2. We devise an adversarial actor–critic reinforcement-learning algorithm that discovers biologically plausible drug–target–disease pathways, yielding interpretable mechanistic explanations for each prediction.
3. We integrate these models within an LLM-driven multi-agent system supporting four practical workflows: (i) disease to drug search or drug to disease search, (ii) pathway elucidation and (iii) automated causal report generation.

Together, these innovations provide a scalable and explainable framework that accelerates the generation and validation of drug repurposing hypotheses while deepening understanding of their modes of action.

## 2 Results

### 2.1 Overview of BioScientist Agent system

The overall structure of the BioScientist Agent is illustrated in Fig. 1. Our objective is to construct an Agent system based on a knowledge graph that encompasses drug repurposing, the discovery of drug mechanisms of action (MoA), and the collection and interpretation of literature related to drug action pathways. Fig. 1 presents the entire workflow of the system:

**Figure 1:**
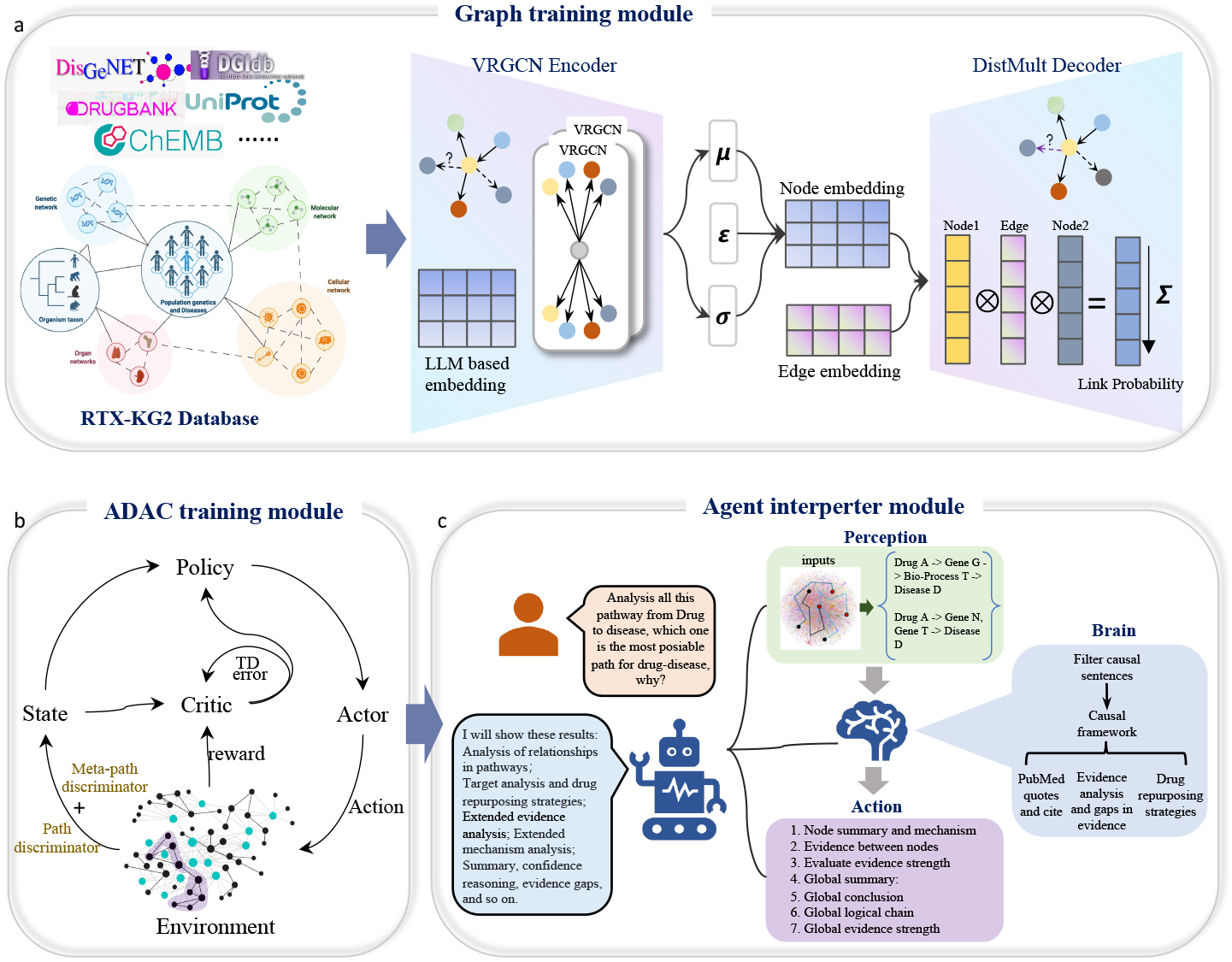
Overview of the BioScientist Agent System. Parts (a), (b), and (c) represent the system’s three modules. (a) Demonstration of RTX-KG2 Knowledge Graph Components and Pretraining with VGAE. This is the Graph Training Module. The left part shows the RTX-KG2 database, which integrates information from sources like DisGeNet, DGIdb, Drugbank, UniProt, and Chembl into a large biological knowledge graph. The graph primarily includes submodules focused on population genetics and diseases, genetic networks, molecular networks, organ networks, cellular networks, and organism taxons. The right part illustrates the pretraining process of the knowledge graph using an encoder-decoder architecture. The model accepts initial features extracted by LLMs for pretraining. The encoder uses a VRGCN model to update nodes, while the decoder uses DistMult to predict edge connections. (b) Adversarial Actor–Critic (ADAC) Model Framework for Drug-Disease MoA Path Extraction. This is the ADAC Training Module. The model receives pre-trained node features, training the actor to generate pathways from drugs to diseases. It also trains a meta-path discriminator and a discriminator to constrain the length and type of these paths. Ultimately, it constructs drug-to-disease MoA pathways and ranks them. (c) LLM-Driven Multi-Agent System Overview. This is the Agent Interpreter Module, which interprets and re-scores MoA pathways generated by the ADAC Training Module. Here, humans request the agent to analyze all pathways from drug to disease. The agent generates analyses of relationships in pathways, target analysis and drug repurposing strategies, extended evidence analysis, extended mechanism analysis, summaries, confidence reasoning, evidence gaps, and more. Internally, the system comprises Perception, Brain, and Action components, responsible for acquiring pathways and related papers, analyzing paper content and pathway logic reliability, and organizing and outputting results.

The first part is the Graph Training Module. Fig. 1a shows the data sources for RTX-KG2, the main connection block, and the graph pre-training process using Variational Relational Graph Convolutional Networks (VRGCN) as the encoder and DistMult as the decoder with RTX-KG2. Each node from RTX-KG2 is provided with its ID, name, synonyms, node type, and description. Before pre-training the knowledge graph, we use Meta’s large language model Llama3.1 and the text to feature representation tool LLM2Vec to represent the textual description of the nodes, obtaining a 4096-dimensional embedding. This is then reduced to 128 dimensions using Gaussian Random Projection, serving as the initial features for the nodes. In the pre-training process for graph representation learning, traditional GCNs mainly focus on single-relation graphs, whereas RGCNs can directly handle graphs containing multiple types of relations [CHG^+^19]. For each edge type (i.e., each semantic relationship) in the graph, an RGCN designs or learns an independent transformation matrix to capture the interactions between nodes under different semantic contexts. Building on this, VRGCN incorporates the concept of a VAE encoder to perform probabilistic modeling of node representations. First, the RGCN propagates messages and aggregates features for each node according to its respective edge types, generating intermediate representations that capture various relational information. Subsequently, for each node the corresponding variational distribution parameters, namely the mean and variance, are computed to construct an approximate posterior distribution. Once the node representations are obtained, the DistMult [YYH^+^14] algorithm is employed to predict the existence of links between nodes. By comparing the predicted links with the actual connections, the model gradually updates and improves its ability to represent the nodes.

The second part is the Adversarial Actor-Critic (ADAC) training module. Fig. 1b shows the ADAC Model Framework for Drug-Disease MoA Path Extraction. After updating the knowledge graph node features using the GVAE pre-training model, we apply a reinforcement learning model ADAC to learn the correct pathways from drugs to diseases [ZWZ^+^20]. The Actor is used for path generation, the Critic evaluates the drug-to-disease effects, and the Meta-path discriminator and path discriminator constrain and assess the types of intermediate nodes and the length of the paths. By using the ADAC algorithm, the generated paths start from the drug, traverse proteins, genes, and pathways, and ultimately reach the correct disease, creating and ranking mechanisms of action (MoA) pathways from drugs to diseases.

The third part is the Agent interperter module. The LLM-Driven Multi-Agent System shows in Fig. 1c. After identifying plausible pathways, their interpretation and experimental validation are conducted through the literature agent system, which assesses the rationality of each path. The system comprises of Perception, Brain, and Action components. Perception is responsible for acquiring pathway information from the ADAC module. The Brain analyzes these pathways, first using an agentic system to filter all related PubMed abstracts for relevant causal sentences then analyzing these sentences using a predefined causal inference framework. The Action component generates the final research reports that include entity summaries and mechanisms, entity-entity relationship analysis, supporting and contradictory evidence analysis, drug repurposing strategies, and pathway chain-of-thought reasoning. Through comprehensive system analysis, for a given drug, disease, and target, the system can obtain extensive information necessary for drug discovery and repurposing, including node summaries and mechanisms, evidence between nodes, evaluation of evidence strength, global summaries, global conclusions, global logical chains, and global evidence strength.

### 2.2 Data Set for BioScientist Agent system

We used RTX-KG2 as the foundational knowledge graph for constructing the BioScientist agent system. The version of RTX-KG2 is v2.9.2, which includes 6,381,804 nodes across 38 distinct categories and 40,989,410 edges across 65 distinct types. Compared to other knowledge graphs, RTX-KG2 contains highly comprehensive information for biological research and drug mechanism discovery, including data on drugs, diseases, genes, and functional experiments results. Additionally, each node and edge is accompanied by extensive literature information for data mining and logical reasoning, which is not available in graphs such as CP-KG or BioKG. CP-KG is composed of 1,246,726 associations between 61,146 entities, and BioKG has 105,465 nodes and 2,043,846 edges, significantly fewer than RTX-KG2. Furthermore, larger knowledge graphs like CKG, which currently comprise close to 20 million nodes and 220 million relationships, include a substantial amount of mutation information. The inclusion of extensive mutation data increases the difficulty of knowledge graph training and representation, and the associations between mutations and phenotypes often result in a high number of false positives. Therefore, taking these factors into account comprehensively, we chose RTX-KG2 for the design of the agent and drug discovery. For RTX-KG2, the top three categories of nodes are ‘Small-Molecular’, ‘OrganismTaxon’ and ‘Gene’ and the top three categories of edges are ‘has participant’, ‘physically interaction with’ and ‘occurs in’ (Fig. 2a). The heatmap of the log values of edge counts illustrates the distribution of node relationships (Fig. 2b). The top five relationships between nodes are concentrated in SmallMolecule to SmallMolecular (2,785,767), Pathway to CellularComponent (2,535,061), Pathway to SmallMolecular (2,268,235), Gene to Disease (1,837,651), and Pathway to OrganismTaxon (1,562,060).

**Figure 2:**
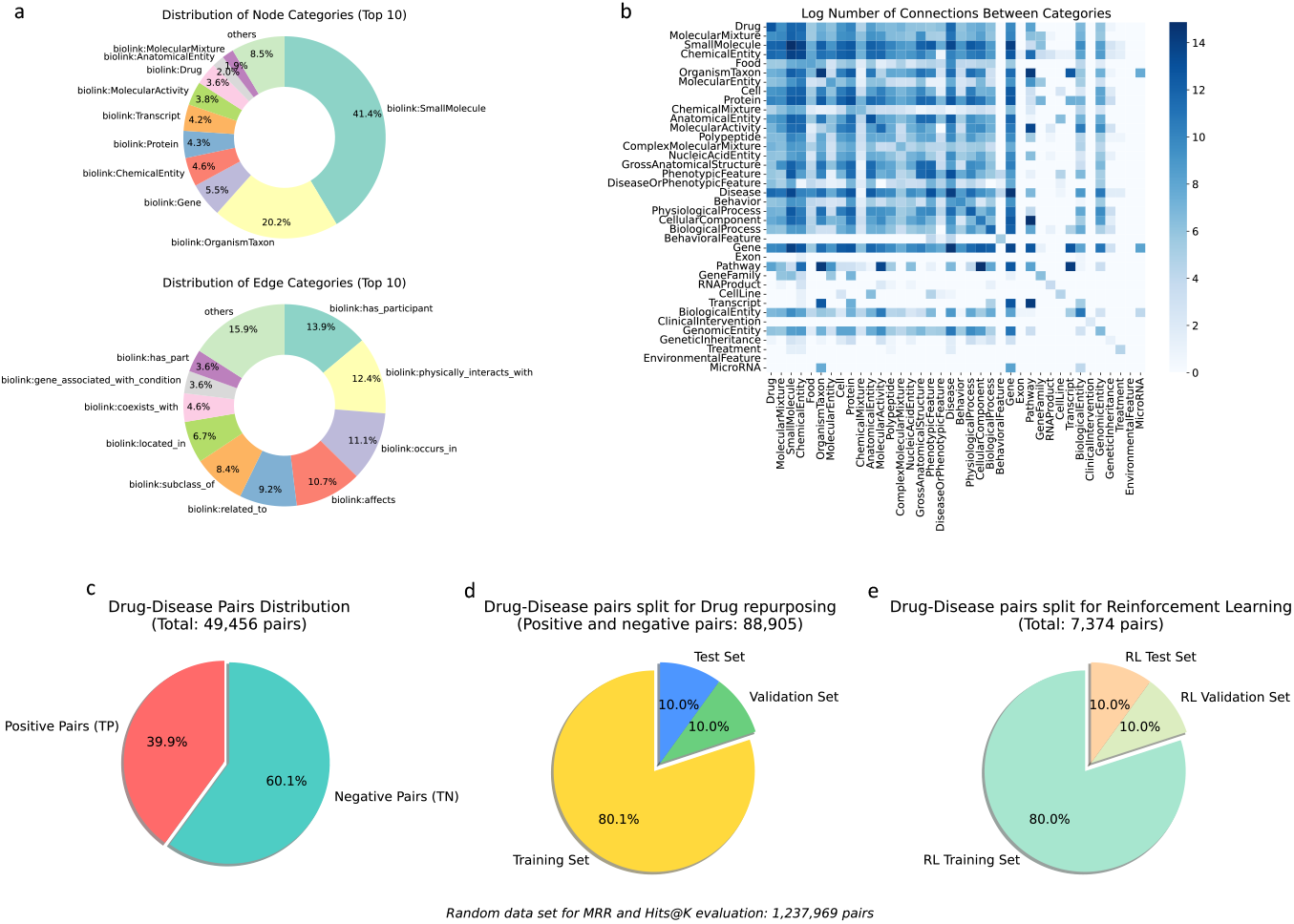
(a) Percentage of nodes by category and percentage of edges by predicate in the customized BKG. (b) Heatmap of log-transformed connection counts between node types. (c) Pie chart of the distribution of true positive and true negative drug-disease relationships. (c) Pie chart of the drug repurposing training dataset distribution. (d) Pie chart of the reinforcement learning training dataset

We obtained real drug-disease relationship data from four databases: MyChemData, SemMedDB Data, NDF-RT Data, and RepoDB Data. This dataset includes positive samples (effective treatments) and negative samples (ineffective treatments). The positive samples (TP pairs) consist of 19,755 known drug-disease treatment relationships, while the negative samples (TN pairs) consist of 29,701 ineffective relationships, totaling 49,456 annotated drug-disease relationships. The distribution of this data is illustrated in Fig. 2c. The training dataset for drug repurposing was divided after feature extraction following pre-training on the knowledge graph, comprising 71,180 pairs for training, 8,857 pairs for validation, and 8,868 pairs for testing. Additionally, there is a random negative sample set of 1,237,969 pairs used for MRR and Hits@K evaluation, with the data distribution shown in Fig. 2d. Furthermore, the reinforcement learning training dataset includes 5,898 expert demonstration drug-disease pairs, 738 pairs for validation, and 738 pairs for testing, with the distribution depicted in Fig. 2e. Based on the classification of drug-disease pairs, approximately 1.88 million four-hop reasoning paths were generated, with 1.51 million in the training set, 0.19 million in the validation and test set, respectively.

### 2.3 BioScientist Agent enables accurate drug repurposing and MoA path predictions

In this study, we evaluated the performance of the BioScientist Agent in drug repurposing and Mechanism of Action (MoA) pathway prediction tasks compared to other models. For drug repurposing, the evaluation metrics included accuracy, F1 score, Mean Reciprocal Rank (MRR), and Hit@K, while MoA pathway prediction utilized MRR and Hit@K. MRR and Hits@K serve as core evaluation metrics, designed based on ranking learning paradigms. In the drug repurposing task, for each true drug-disease therapeutic relationship, a candidate set, comprising the true pair and related negative samples (random data set from Fig. 2d), was constructed. The model predicted probabilities to rank the candidate set in descending order, subsequently calculating the position of the true pair or MoA pathway within the ranking list. Negative samples in drug repurposing were constructed using a dual matching strategy, collecting random pairs with the same drug and same disease as distractors. MRR assesses overall ranking quality by calculating the average reciprocal rank of the true pair across all queries, with a range of [0,1], where higher values indicate the model ranks true therapeutic relationships higher. Hits@K measures the proportion of true pairs appearing in the top K candidates, providing a more intuitive top-K recommendation performance metric. In MoA pathway prediction tasks, a dataset containing all paths was constructed, excluding paths from the training and validation sets. Paths were scored and ranked, observing the ranking of the correct path and calculating MRR and top-K metrics. These indicators effectively reflect the BioScientist Agent’s capability in its drug repurposing algorithm and MoA pathway search within real application scenarios.

In the drug repurposing task, compared to existing high performance algorithms, including Graph-Sage, GAT, TAGConv, GCN, TransformerConv, and SGFormer, the BioScientist Agent utilizing VGAE demonstrated superior performance, improving accuracy and F1 score by approximately 3.3% to 3.7%. Key metrics such as MRR, Hit@1, and Hit@3 improved by approximately 36.32%, 52.76%, and 42.34% over the second-place TransformerConv. In MoA pathway prediction tasks, the BioScientist Agent using pretrained node features also exhibited enhanced performance, surpassing KGML-xDTD by approximately 11.01%, 15.25%, and 5.18% in MRR, Hit@1, and Hit@10, respectively. The complete results for drug repurposing and MoA pathway prediction are presented in Tables 1 and 2. The strong performance of the BioScientist Agent in drug repurposing and MoA pathway prediction provides a solid foundation for downstream pathway interpretation.

**Table 1:**
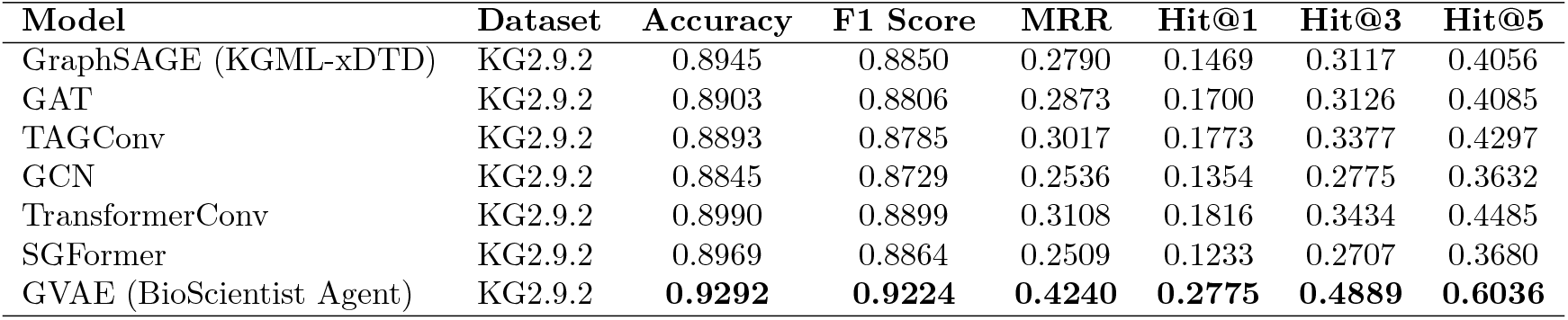
Performance Metrics for Drug Repurposing Across Different Models Using Dataset KG2.9.2.

| Model | Dataset | Accuracy | F1 Score | MRR | Hit@1 | Hit@3 | Hit@5 |
| --- | --- | --- | --- | --- | --- | --- | --- |
| GraphSAGE (KGML-xDTD) | KG2.9.2 | 0.8945 | 0.8850 | 0.2790 | 0.1469 | 0.3117 | 0.4056 |
| GAT | KG2.9.2 | 0.8903 | 0.8806 | 0.2873 | 0.1700 | 0.3126 | 0.4085 |
| TAGConv | KG2.9.2 | 0.8893 | 0.8785 | 0.3017 | 0.1773 | 0.3377 | 0.4297 |
| GCN | KG2.9.2 | 0.8845 | 0.8729 | 0.2536 | 0.1354 | 0.2775 | 0.3632 |
| TransformerConv | KG2.9.2 | 0.8990 | 0.8899 | 0.3108 | 0.1816 | 0.3434 | 0.4485 |
| SGFormer | KG2.9.2 | 0.8969 | 0.8864 | 0.2509 | 0.1233 | 0.2707 | 0.3680 |
| GVAE (BioScientist Agent) | KG2.9.2 | **0.9292** | **0.9224** | **0.4240** | **0.2775** | **0.4889** | **0.6036** |

**Table 2:** Performance Metrics for MoA path predictions between KGML-xDTD and Bioscientist Agent.

|  | MRR | Hit@1 | Hit@10 | Hit@50 | Hit@100 | Hit@500 |
| --- | --- | --- | --- | --- | --- | --- |
| KGML-xDTD | 0.109 | 0.059 | 0.193 | 0.496 | 0.613 | 0.849 |
| BioScientist agent | 0.121 | 0.068 | 0.203 | 0.508 | 0.619 | 0.864 |

### 2.4 BioScientist Agent elucidating RL-based pathway and underlying mechanisms

We integrated an LLM-based causal agent to evaluate the proposed drug-disease relationships and corresponding MoA. The ADAC model may propose thousands of potential drug-disease pathways and intermediary drug targets for any given disease, so being able to elucidate the underlying mechanisms and biological plausability of each would prove useful for ranking and identifing the most salient ones for further research. For each entity-entity relationship, we queried PubMed abstracts for entity co-occurrences using PubMed Entrez to find semantically related articles. The abstracts were then filtered using an agent-based approach to find unique causal sentences, which were used as foundational evidence for general-purpose GPT-4 models to analyze using a provided causal inference framework. The final output is a list of relevant PubMed quotations used by the LLM in its analaysis, evidence evaluation, and a final confidence score *C* ∈ [0, 1] to indicate the causal strength between the two entities. We obtained the overall score for a pathway by getting the geometric mean of all the scores 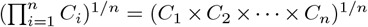 where n is the number of biological entities in the path. Using this pipeline of evidence-based scoring and causal frameworks overcomes common limitations of LLM-based approaches by minimizing speculative reasoning and anchoring all conclusions in peer-reviewed literature. We show more details regarding causal sentences and the causal inference framework used in Section 4.2.3. On a test run with 40 selected pathways, 20 scientifically-tested positive pathways and 20 randomly chosen negative pathways, our agent achieved an ROC AUC of 0.8562, with an accuracy of 0.8750, F1-score 0.8837, precision 0.8261, and recall 0.9500. The higher false positive rate is due to the random selection of negative pathways. Compared to adversarial actor–critic (ADAC) reinforcement learning model, which had an ROC AUC of 0.642, accuracy of 0.539, F1-score of 0.679, and precision of 0.529, the BioScientist Agent performed significantly better. Fig. 3 illustrates these performance metrics, highlighting the superior results of the BioScientist Agent in all areas except recall, where both models were equal.

**Figure 3:**
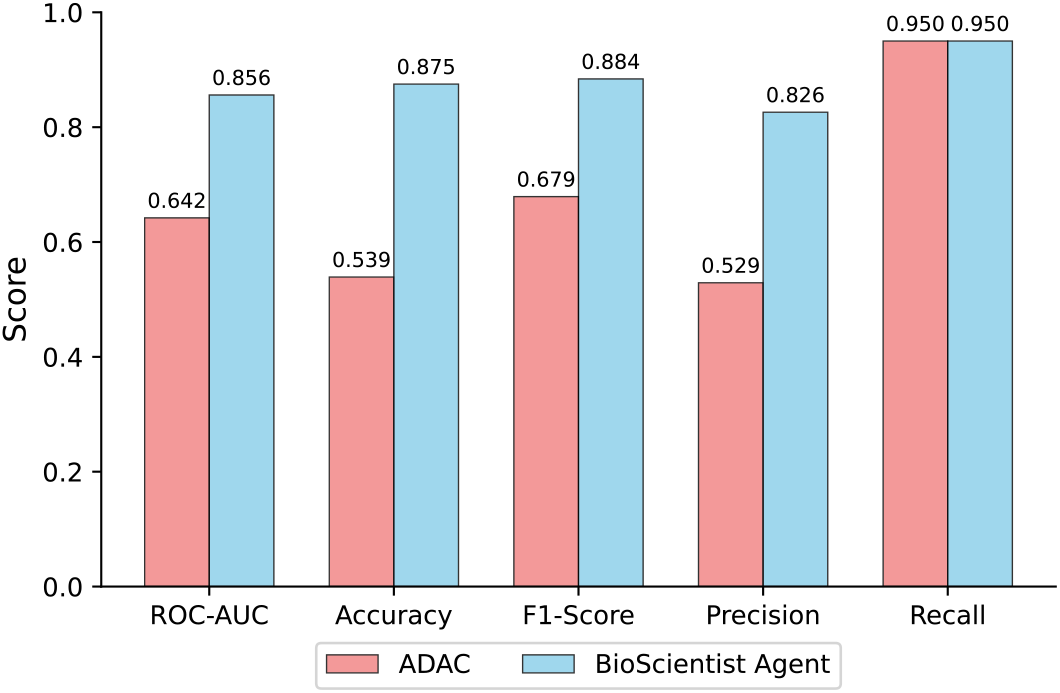
Path classification performance comparison between adversarial actor–critic (ADAC) and BioScientist in 40 selected pathways.

### 2.5 BioScientist Agent Mining Target Information

Even without invoking any knowledge graph pre-training models or reinforcement learning models, our system enables information mining based on knowledge graph data. This example demonstrates how we leverage the capabilities of knowledge graphs to identify mechanisms of action (MoA) associated with the targets Protein arginine methyltransferase 5 (*PRMT5*) and Fascin actin-bundling protein 1 (*FSCN1*). Using these graphs, we can quickly access relevant MoA pathways for *PRMT5* and *FSCN1*, as illustrated in Fig. 4. This enables us to rapidly gather all potential biological concept nodes related to the targets, with a focus on drug molecules, associated proteins, and diseases. Additionally, because each node pair has related PubMed IDs, we can utilize APIs to obtain the impact factor distribution of all papers within a pathway, allowing us to rank and analyze the pathway’s reliability. This helps researchers pinpoint trustworthy papers for scientific study and discovery (Fig. 4). Our findings reveal that the top three most plausible pathways associated with *PRMT5* are: carbamazepine - infectious disease - *PRMT5* - epilepsy; dexamethasone - *PRMT5* - CRK - atopic dermatitis; nitroglycerin - bone morphogenetic protein 2 - *PRMT5* - angina pectoris. Several related papers can indirectly demonstrate the reliability of the pathways. For instance, Zhang et al. identified *PRMT5* as a gene associated with epilepsy [ZLW^+^24]. Kim et al. emphasized the significant role of *PRMT* family genes, especially *CARM1, PRMT1, PRMT5*, and *PRMT6*, in regulating inflammatory responses, particularly in modulating inflammation-related transcription factors or cofactors [KYY^+^16]. The top three pathways linked to *FSCN1* are: gefitinib - cell proliferation - *FSCN1* - non-small cell lung carcinoma; docetaxel - *MAPT* - *FSCN1* - breast neoplasm; capecitabine - *GNAS* - *FSCN1* - colonic neoplasm. In the study by Luo et al., the association between *FSCN1* protein expression and clinicopathological features, including survival rates, was explored in 156 non-small cell lung cancer (NSCLC) patients. It was found that *FSCN1* mRNA and protein expression levels were significantly higher in NSCLC tissues compared to normal lung tissues. Additionally, elevated *FSCN1* protein expression was significantly associated with poorer overall survival rates in NSCLC patients. Multivariate analysis indicated that increased *FSCN1* protein expression is an adverse prognostic biomarker for NSCLC patients [LYL^+^15]. The study by Alajez et al. demonstrated that the combined overexpression of both BMI1 and *FSCN1* may serve as a superior prognostic marker for colonic neoplasm compared to either marker alone. Targeting BMI1 and *FSCN1* may also offer potential therapeutic opportunities in colonic neoplasm [Ala16].

**Figure 4:**
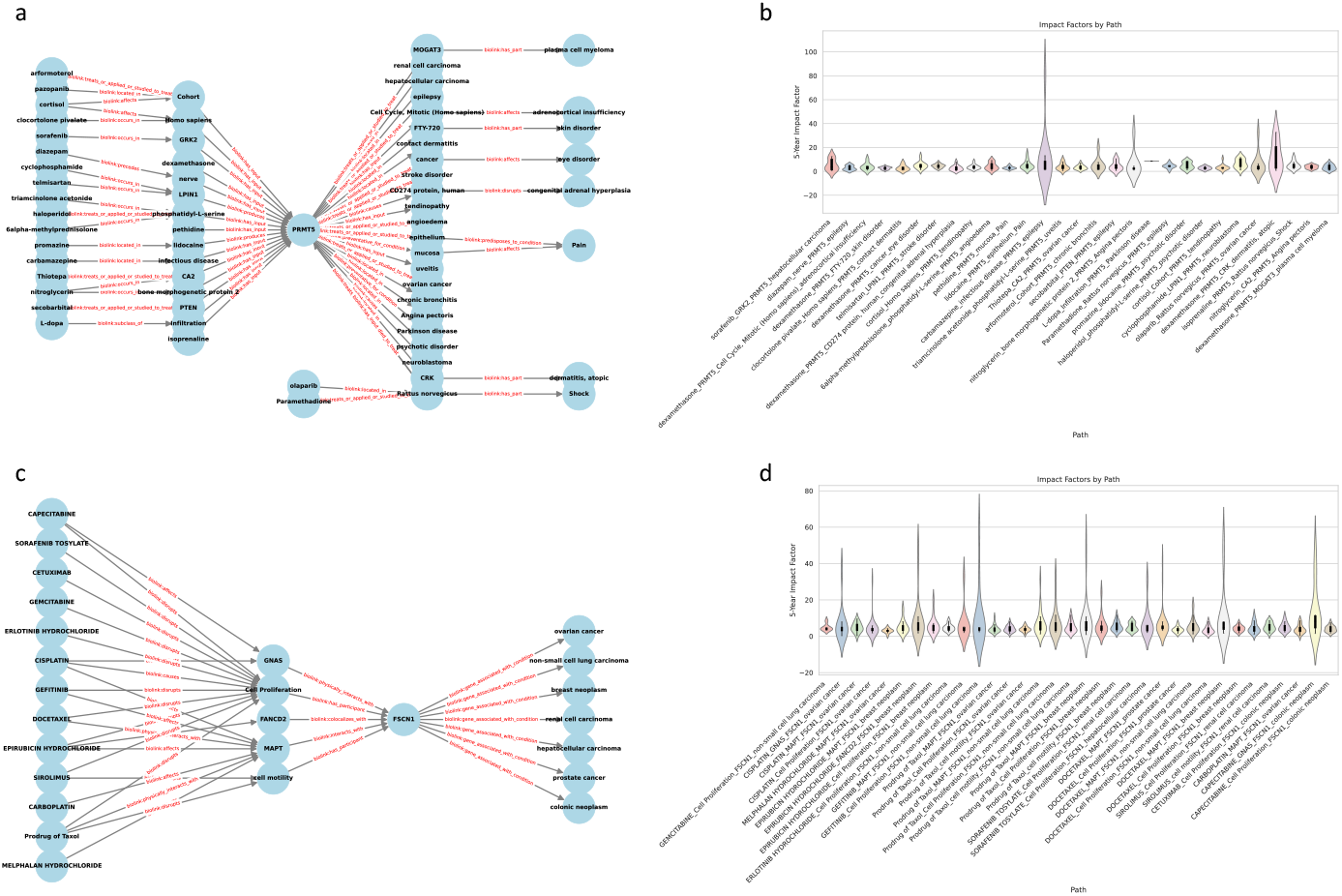
MoA pathway profiles centered on targets PRMT5 and FSCN1, and the distribution of impact factors of all publications involved in the pathways: (a) Extraction of the MoA pathway centered on target *PRMT5*; (b) Violin plots showing the distribution of impact factors for all publications between every pair of nodes in each *PRMT5*-associated pathway; (c) Extraction of the MoA pathway centered on target *FSCN1*; (d) Violin plots showing the distribution of impact factors for all publications between every pair of nodes in each *FSCN1*-associated pathway.

### 2.6 BioScientist Agents interpret MoA pathways

The BioScinetsi Agent has the ability to finely interpret an MoA pathway. We use carefully designed prompts for paper information extraction and interpretation to score the reliability of the MoA pathway. The prompt for the multi-agent approach to extract causal information from PubMed abstracts is shown in Supplementary Material 1. Each agent analyzes a provided list of PMIDs and corresponding abstracts to extract either supporting or contradictory sentences and returns that information. In Supplementary Material 2, we show the causal analysis framework used to determine the causal relationship between two entities, including mechanistic evidence, association strength, biological plausibility, literature consistency, causal relationship analysis, and overall confidence. The global summary of a drug-gene pathway provided by the agent is shown in Material 3. This includes a summary of the most notable evidence in determining the causal score along with the PMID citations for the evidence. The agent evaluates both supporting and contradictory evidence to identify gaps that limit the pathway’s score. The final output is a JSON file, with the overall pathway analysis part shown in Fig. 5. The complete JSON file is in Supplementary Material 4.

**Figure 5:**
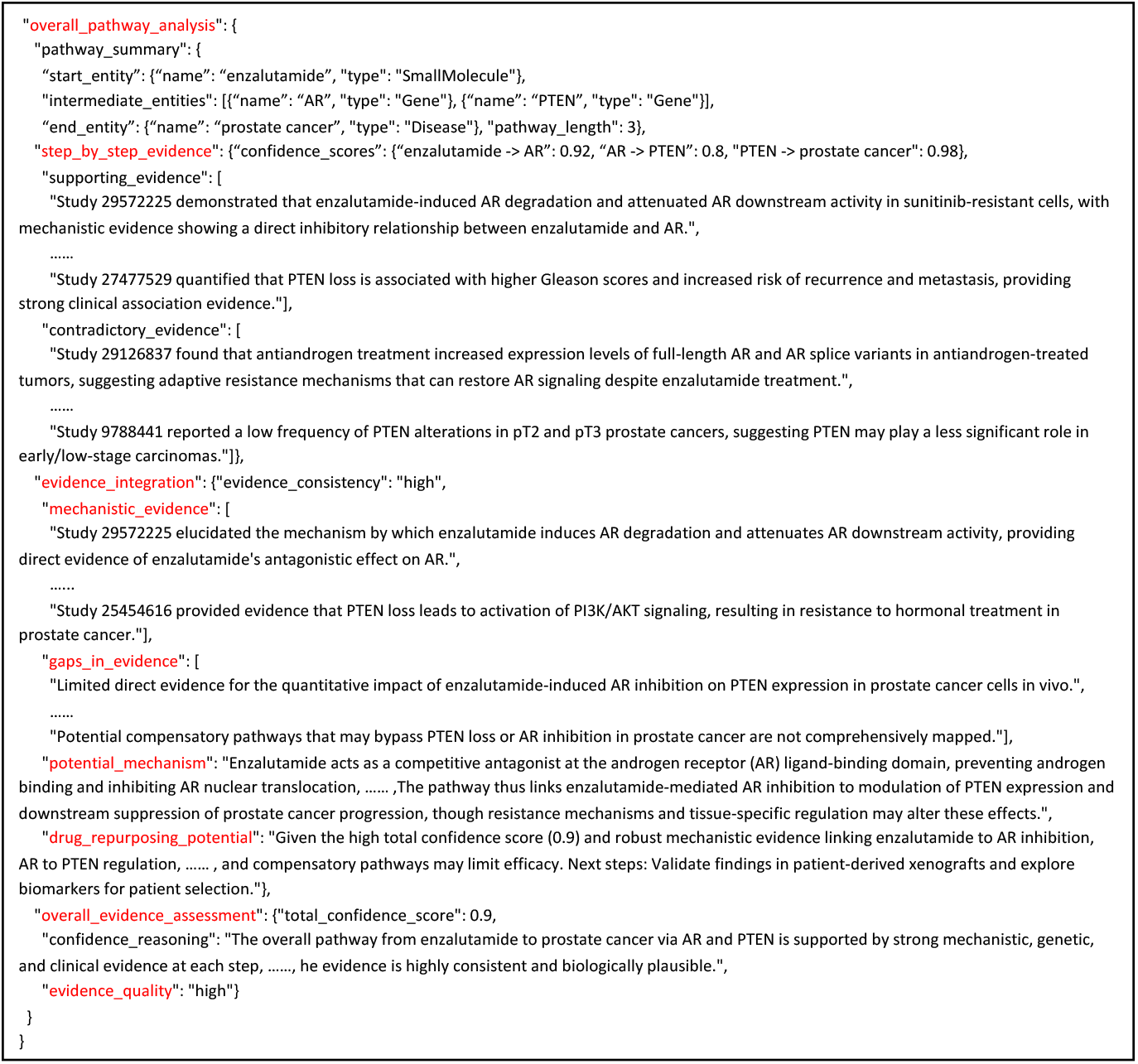
Demonstration of the overall pathway analysis in MoA pathway interpretation, with complete results in Supplementary Material 4.

In the JSON file, we analysed the putative repurposing of enzalutamide (Xtandi) for mitigating prostate cancer through the targets of the *AR* gene and the *PTEN* gene. The agent produced a causal confidence score of C=0.9 based on evidence extracted from 1025 articles. Each individaul entity-entity relationship within the path has its own analysis of BH criteria, mechanistic evidence, causal terminology, potential mechanism, key strengths/weaknesses, and citations. Key causal sentences used in the agent analysis were identified and then summarized in the global summary. For example: “Study [ PMID: 23701654][DSS^+^13] demonstrated that enzalutamide acts as an androgen receptor antagonist, inhibiting AR signaling pathways, supporting a strong causal relationship between enzalutamide and *AR*.”, “Study [30177856] [dLPRDC^+^18] showed that enzalutamide treatment is associated with reduced AR activity, indicating a direct effect of enzalutamide on AR levels.”, “Study [PMID: 21532617] [WRH^+^11] elucidated that AR inhibits *PTEN* transcription in prostate cancer cells, establishing a causal link between AR and *PTEN*.”, “Study [PMID: 23902739] [CLSK^+^13] reported that loss of *PTEN* is frequently observed in cancer, resulting in the deregulation of cell survival, growth, and proliferation, indicating a strong association between *PTEN* loss and prostate cancer progression.” The agent also proposed a potential pathway MoA based on the mechanistic evidence it identified (“The potential mechanism involves enzalutamide binding to the androgen receptor (AR), preventing its activation and subsequent transcription of target genes. This inhibition leads to decreased *PTEN* expression, as AR negatively regulates *PTEN* transcription. The loss of *PTEN* function activates the *PI3K* /*AKT* signaling pathway, promoting cell survival and proliferation, ultimately contributing to prostate cancer progression.”). For high-confidence paths, the agent proposes detailed drug repurposing strategies based on the evidence provided: “Given the total confidence score of 0.9, a candidate drug for repurposing could be a selective AR modulator (SARM) that has shown promise in modulating AR activity without the resistance seen with enzalutamide. The experimental design would involve using prostate cancer cell lines with varying levels of AR expression and *PTEN* status. Controls would include untreated cells and cells treated with enzalutamide. Endpoints would include measuring AR and *PTEN* expression levels through qPCR and Western blotting, as well as assessing cell proliferation and apoptosis through MTT assays and flow cytometry. The experiment would test the candidate drug’s effect on the pathway by evaluating whether it can effectively reduce AR activity and restore *PTEN* levels compared to enzalutamide. Expected outcomes would include a reduction in AR activity and an increase in *PTEN* levels, supporting the hypothesis that the candidate drug could be effective in overcoming resistance mechanisms. Potential challenges include variability in patient-derived cell lines and the need for in vivo validation. Next steps would involve optimizing the dosing regimen and exploring combination therapies with existing treatments.” Identified weak links such as “Further research is needed to explore the specific mechanisms by which AR variants contribute to enzalutamide resistance.”, and “More studies are required to clarify the role of other signaling pathways that may interact with the *AR-PTEN* axis in prostate cancer” were flagged for further wetlab investigation. Collectively, this framework provides viable and substantiated biological causal pathways that are traceable to primary literature by combining epidemiological causal inference and machine reasoning to deliver a transparent and rigorous solution.

## 3 Discussion

Knowledge graphs encompass an extensive amount of information, and leveraging them can accelerate tasks such as drug repurposing and predicting drug mechanisms of action. This not only speeds up the progress of drug development but also reduces research and development costs. In this study, we developed the BioScientist Agent system, which integrates graph neural network algorithms, ADAC reinforcement learning algorithms, and an agent-based literature interpretation and path scoring system. This comprehensive system manages the entire workflow, including knowledge graph training, edge discrimination, MoA path searching, and result interpretation, thereby playing a crucial role in accelerating drug development.

Previous research seldom involved training on such large-scale knowledge graphs due to computational constraints. Typically, these studies limited the graphs to 4-5 types of nodes, such as drugs, diseases, and genes, encompassing tens of thousands of nodes and millions of edges. They excluded additional node types like cell types, transcription factors, and microRNA, as well as numerous edges supported by only a few publications. In contrast, our study incorporates all edges deemed present in existing research into the training scope, with the number of nodes reaching the millions and the number of edges reaching tens of millions. This results in a highly comprehensive dataset, significantly enhancing the model’s ability to discover new and potential drug-disease relationships and to explore drug mechanisms of action, even for those relationships with minimal prior research.

Thanks to continuous advancements in biological science research, the data sources for knowledge graphs are constantly being updated. We believe that an Agent system capable of rapid iteration is of greater significance. Currently, the BioScientist Agent can quickly iterate when handling updated databases; however, it still requires retraining from scratch to associate new nodes with existing ones using graph neural network algorithms. This process is time-consuming and necessitates extensive data preprocessing and initial feature extraction for nodes. In our future work, we aim to establish mechanisms for the rapid integration of new nodes and edge relationships. This includes updating nodes through local graph representation capabilities and incorporating more node representation models to enhance their representation abilities. For instance, we plan to utilize large model tools like ‘SMI-TED289M’ that encode molecular SMILES for feature representation [SSB^+^24]. By training on drug molecules that have some experimental data but still possess many unknown functions, we aim to predict targeted diseases and the mechanisms of action (MoA) of these unknown molecules. Ultimately, this will enable the BioScientist Agent system to exert a greater impact.

## 4 Methods

### 4.1 Datasets

#### 1. Original datasets

We used the canonicalized version of Reasoning Tool X Knowledge Graph 2 (RTX-KG2c) v2.9.2 for constructing, training, and extracting drug-disease pathways from the knowledge graph. It integrates a large amount of human-curated and publication-based database knowledge and categorizes nodes and edges based on the Biolink model. The RTX-KG2c version v2.9.2 we used contains approximately 6,873,228 nodes and 46,568,039 edges, covering knowledge from 70 public biomedical sources, with all biological concepts represented as nodes and all concept-predicate-concept relationships represented as edges. After excluding node types unrelated to biological explanations and edges with redundant concepts, the graph retains 6,381,804 nodes with 37 distinct categories (Fig. 1A) and 40,989,410 edges with 67 distinct types (Fig. 1B) for downstream model training.

#### 2. Demonstration paths

Demonstration paths are prepared before conducting reinforcement learning because drugs treat diseases through specific targets. We constructed certain drug-to-disease paths, such as drug–gene–protein–disease and drug–drug–protein–disease. We extracted 1.9 million demonstration paths from the knowledge graph, based on known drug-target interactions collected from two curated biomedical data sources: DrugBank (v5.1) and Molecular Data Provider (v1.2), as well as the normalized Google distance (NGD) score based on PubMed publications, with NGD scores required to be less than 0.6. The PubMed publication-based NGD [53] defined below:

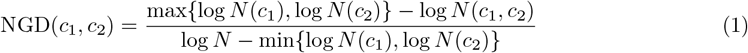

where *c*_1_ and *c*_2_ are two biological concepts used in the customized BKG, *N* (*c*_1_) and *N* (*c*_2_) respectively represent the total number of unique PubMed IDs associated with *c*_1_ and *c*_2_, *N* (*c*_1_, *c*_2_) is the total number of unique PubMed IDs shared between *c*_1_ and *c*_2_, and *N* is the total number of pairs of Medical Subject Heading (MeSH) terms annotations in the PubMed database.

### 4.2 Models

The framework of BioScientist Agent consists of 3 modules: a VGAE-based Graph Neural Network Representation Model [KW16] for drug repurposing, an adversarial actor-critic RL model for drug-to-disease pathway scoring and a proposed LLM-powered multi-agent system for automated scientific discover and MoA path re-scoring and interpreting. The architecture of the whole Agent system showed in Fig2.

#### 4.2.1 VGAE-based Graph Neural Network Representation Model

##### Model Architecutre

Let *G* = (*V, E*) be a directed biomedical knowledge graph, where:

- Each node *v* ∈ *V* represents a biological entity (e.g., a specific drug, disease, gene, or pathway)
- Each edge *e* ∈ *E* represents a biomedical relationship

Given *G*, we employ a VGAE to learn low-dimensional node embeddings **Z** ∈ ℝ^|*V* |*×d*^, where *d*≪ | *V*|. The model consists of the following components:

###### 1. Variational Relational Graph Convolutional Network (VRGCN) Encoder

The encoder uses a two-layer relational graph convolutional network with separate variational heads to infer stochastic embeddings:

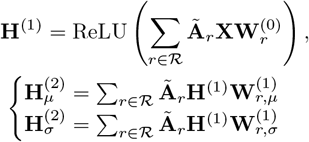

where:

- 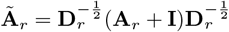 is the normalized adjacency matrix for relation *r* ∈ *ℛ* with self-loops
- **X** ∈ ℝ^|*V*|*×m*^ is the node feature matrix with *m*-dimensional features
- 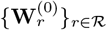 are first-layer relation-specific weight matrices
- 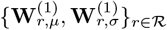 are second-layer weight matrices for mean and variance heads
- ReLU(*x*) = max(0, *x*) is the rectified linear unit activation function
- *ℛ* is the set of relation types in the graph

The latent embeddings **Z** are sampled using the reparameterization trick:

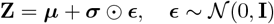

where:

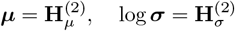

- Separate RGCN heads compute ***µ*** and log ***σ*** to enable variational learning
- The architecture supports multiple relation types through relation-specific parameters
- Dropout is typically applied between layers for regularization

###### 2. Decoder for Link Prediction

The decoder uses a DistMult approach for link prediction by reconstructing edges:

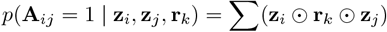

where:

- **z**_*i*_ and **z**_*j*_ are the source and destination node embeddings
- **r**_*k*_ is the relation embedding for the edge type
- ⊙ denotes element-wise multiplication

The reconstructed adjacency matrix **Â** is derived from these scores for each relation type. The parameters are initialized using Xavier uniform initialization to ensure proper scaling of inputs and outputs.

##### Model Training

The model optimizes the loss function that combines reconstruction and regularization:

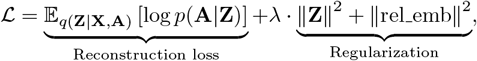

where:

- The reconstruction loss is computed using binary cross-entropy with logits.
- Regularization is applied to both node embeddings **Z** and relation embeddings rel emb.
- *λ* controls the regularization strength (default: *λ* = 0.01).

##### Model Evaluation

Our evaluation pipeline includes several key components designed to assess the performance of the VGAE model on knowledge graph link prediction tasks. We first obtain true drug-disease interactions, including true positives and true negatives, from MyChemData, SemMedDB Data, NDF-RT Data, and RepoDB Data. Positive samples (known drug-disease treatment relationships) are divided into two subsets: expert demonstration pairs based on knowledge graph paths and other positive pairs, ensuring effective utilization of high-quality samples. During the division process, a drug-centered hierarchical sampling strategy is adopted. Single-indication drugs are prioritized for the training set, while multi-indication drugs are stratified using drug IDs to prevent data leakage by dispersing different indications of the same drug across datasets. We designed a balanced random negative sample generation mechanism, creating random pairings for each drug and disease in the positive samples, ensuring they do not duplicate known positives or negatives. Random negative samples are labeled as an independent category (label 2) and are mainly used to support ranking evaluation metrics like MRR (Mean Reciprocal Rank) and Hits@K in link prediction tasks. The final dataset composition for each set (training/validation/testing) includes expert demonstration positives, other positives, true negatives, and random negatives, with random shuffling ensuring uniform sample distribution. Additionally, a large-scale set of random pairs is generated specifically for ranking performance evaluation in the validation and testing phases, providing a comprehensive and rigorous evaluation benchmark for drug repurposing models.

1. **Embedding Generation and Feature Construction**. The evaluation process begins by loading a pre-trained VGAE model and generating node embeddings: (1) Load the best trained model checkpoint and set it to evaluation mode; (2) Generate embeddings for all nodes using the encoder: *z* = Encoder(*X, E, R*), where *X* represents the init node feature matrix, *E* denotes the edge index matrix defining graph connectivity, and *R* represents the edge type/relation indices for each edge; (3) Create a dictionary mapping node IDs to their corresponding embeddings. For link prediction, we construct feature vectors by concatenating source and target node embeddings: **x**_*pair*_ = [**z**_*source*_ *⊕* **z**_*target*_], where *⊕* denotes concatenation, resulting in 256-dimensional feature vectors.
2. **Classification and Prediction**. A Random Forest classifier is employed as the final prediction layer with 2000 estimators, maximum depth of 35, square root of features per split, and automatic class weight balancing to handle imbalanced datasets. The classifier is fitted on concatenated embeddings from training pairs.
3. **Evaluation Metrics**. We employ multiple evaluation metrics to comprehensively assess model performance. Classification metrics include: (1) *Accuracy* :

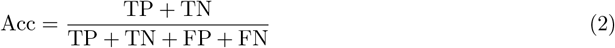

where TP, TN, FP, FN are true positives, true negatives, false positives, and false negatives, respectively; (2) *Macro F1-score*:

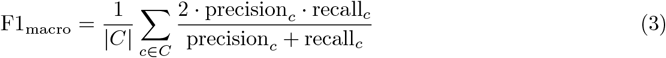

where |*C*| is the number of classes; (3) *Micro F1-score*:

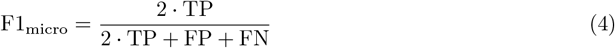

aggregated globally across all classes. Ranking metrics for link prediction evaluation include: (1) *Mean Reciprocal Rank (MRR)*:

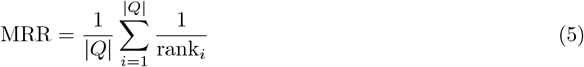

where |*Q*| is the number of positive query pairs and rank_*i*_ is the rank of the correct answer among all candidates for query *i*; (2) *Hit@K* :

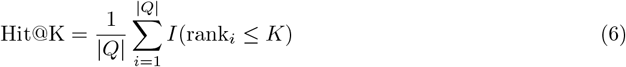

where *I*(·) is the indicator function that equals 1 if the condition is true and 0 otherwise, evaluated for *K* ∈ {1, 3, 5}.

#### 4.2.2 Adversarial Actor-Critic RL Model

We formulate the MOA prediction as a path-finding problem and adapt the adversarial actor-critic reinforcement learning model to solve it. Reinforcement learning is defined as a Markov decision process (MDP) that contains:

##### Basic concept

- **States**: Represent the current situation of the agent within the environment. Each state *s*_*t*_ at time *t* is defined as:

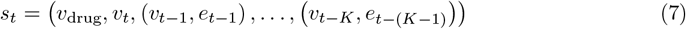

where:
  - *v*_drug_ ∈ *V*_drug_ is the given starting drug node.
  - *v*_*t*_ ∈ *V* represents the node where the agent is located at time *t*.
  - *v*_*t*−*K*_, *e*_*t*−(*K*−1)_ represents the previous *K*th node and the (*K* − 1)th predicate.
  For the initial state *s*_0_, the previous nodes and predicates are substituted by a special dummy node and predicate. The state embedding **s**_*t*_ is obtained by concatenating the embeddings of all nodes and predicates in *s*_*t*_, where:
  - Node embeddings are generated using the LLM2Vec model.
  - Predicate embeddings are represented by one-hot vectors.
- **Actions**: Define the possible moves the agent can make. The action space *A*_*t*_ at each node *v*_*t*_ includes a self-loop action *a*_self_ and actions to reach its outgoing neighbors in the knowledge graph *G*. To manage memory limitations and handle nodes with large out-degrees, neighbor actions are pruned based on PageRank scores if a node has more than 3,000 neighbors. Specifically, the action space is defined as:

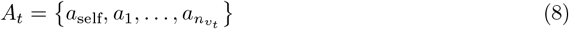

where 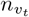 is the out-degree of node *v*_*t*_ ∈ *V*. For each action *a*_*t*_ = (*e*_*t*_, *v*_*t*+1_) ∈ *A*_*t*_ taken at time *t*, its embedding **a**_*t*_ is obtained by concatenating its node and predicate embeddings. Two embedding matrices are learned:

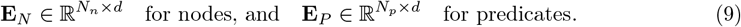

where:
  - *d* is the embedding dimension.
  - *N*_*n*_ is the number of nodes in the graph.
  - *N*_*p*_ is the number of predicate categories.
  - Each subnetwork uses separate embedding matrices.
- **Rewards**: Provide feedback to the agent based on the actions taken. During the path-searching process, the agent only receives a terminal reward *R*_*e*,*T*_ from the environment. There are no intermediate rewards from the environment, i.e.,

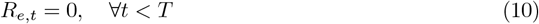

Let *v*_*T*_ be the last node of the path, and *N*_drug_ be the known diseases that drug *v*_drug_ can treat. The terminal reward *R*_*e*,*T*_ from the environment is calculated using the drug repurposing model as follows:

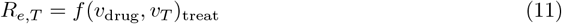

where *p*_treat_ is the “treat” class probability predicted by the drug repurposing model *f*.
- **Policy**: Determines the agent’s behavior by mapping states to actions.

##### Model Architecture

The adversarial actor–critic RL model comprises four subnetworks that share the same Multilayer Perceptron (MLP) architecture MLP_*i*_, where *i* identifies each subnetwork (e.g., *a* for actor, *c* for critic). However, they have different parameters:

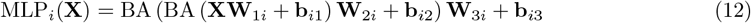

where: {**W**_1*i*_, **W**_2*i*_, **W**_3*i*_, **b**_*i*1_, **b**_*i*2_, **b**_*i*3_} are the parameters and biases of linear transformations. BA represents a batch normalization layer followed by an Exponential Linear Unit (ELU) activation function.

The Model Architecture consists of the following components:

- **Actor Network**: The actor network learns a path-finding policy *π*_*θ*_, where *θ* represents all parameters of the actor network. This policy guides the agent to choose an action *a*_*t*_ from the action space *A*_*t*_ based on the current state *s*_*t*_:

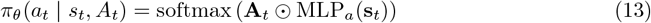

where:
  - **A**_*t*_ is the embedding matrix of the action space *A*_*t*_.
  - ⊙ denotes the dot product.
  *π*_*θ*_(*a*_*t*_ | *s*_*t*_, *A*_*t*_) represents the probability of choosing action *a*_*t*_ at time *t* from the action space *A*_*t*_ given the state *s*_*t*_.
- **Critic Network**: The critic network estimates the expected reward *Q*_*ϕ*_(*s*_*t*_, *a*_*t*_), where *ϕ* represents all parameters of the critic network, if the agent takes action *a*_*t*_ at state *s*_*t*_:

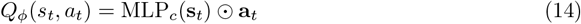
- **Path Discriminator Network**: Since the RL agent only receives a terminal reward *R*_*e*,*T*_ indicating whether it reaches an expected target, a path discriminator network is introduced to encourage the agent to find biologically reasonable paths and provide intermediate rewards. This network acts as a binary classifier distinguishing whether a path segment (*s*_*t*_, *a*_*t*_) is from demonstration paths or generated by the actor network. Positive Samples: Known demonstration path segments 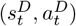. Negative Samples: Actor-generated non-demonstration path segments 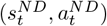. The path discriminator is defined as:

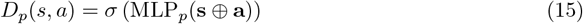

where:
  - *σ* is the sigmoid function.
  - *⊕* denotes concatenation.
  - **s** and **a** are the embeddings of state *s* and action *a*, respectively.
  The discriminator is optimized with the loss function:

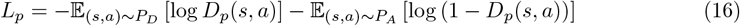

where:
  - *P*_*D*_ is the demonstration path segment distribution.
  - *P*_*A*_ is the actor-generated non-demonstration path segment distribution.
  Based on the discriminator’s output *D*_*p*_(*s*_*t*_, *a*_*t*_), the intermediate reward *R*_*p*,*t*_ is calculated as:

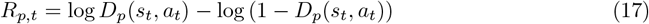
- **Meta-Path Discriminator Network**: Similarly, the meta-path discriminator network judges whether the meta-path of the actor-generated paths is similar to that of demonstration paths. A meta-path is defined as the sequence of node categories in the path (e.g., “Drug” → “Gene” → “Gene” → “Disease”, “Drug” → “Gene” → “Biological Process” → “Disease”). The meta-path discriminator is defined as:

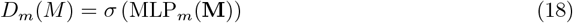

where:
  - **M** is the embedding of the meta-path *M*, obtained by concatenating the learned category embeddings of all nodes in the path.
  It is optimized with the loss function:

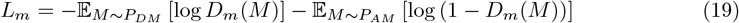

where:
  - *P*_*DM*_ is the distribution of demonstration meta-paths.
  - *P*_*AM*_ is the distribution of actor-generated non-demonstration meta-paths.
  The intermediate reward *R*_*m*,*t*_ from the meta-path discriminator is computed as:

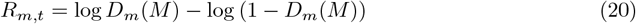
- **Integrated Intermediate Reward**: The integrated intermediate reward *R*_*t*_ at time *t* combines the rewards from both discriminators and the terminal reward:

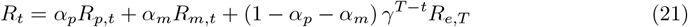

where:
  - *α*_*p*_ ∈ [0, 1] and *α*_*m*_ ∈ [0, 1 − *α*_*p*_] are hyperparameters.
  - *γ* is the decay coefficient.
  - *R*_*e*,*T*_ is the terminal reward from the environment.

##### ADAC Model Training

From the complete library of expert demonstration paths (containing all inference paths from drugs to diseases), precise matching and extraction are performed based on the drug-disease pairs in the pre-divided training, validation, and test sets. Specifically, this involves converting drug-disease pairs into their corresponding entity IDs and then searching the expert path data for all complete inference paths that start with the specified drug and end with the specified disease. This matching process ensures that each data subset contains only the expert demonstration trajectories strictly corresponding to its drug-disease pairs, maintaining data independence across the training, validation, and test sets. Since a single drug-disease pair may have multiple inference paths, the final expert demonstration dataset retains the diversity and completeness of paths.

###### 1. Initialization

- The actor network is initialized using the behavior cloning method [**?**], where the training set of demonstration paths guides the agent’s sampling using a Mean Squared Error (MSE) loss.

###### 2. Training Discriminators

- For the first *z* epochs, the parameters of the actor and critic networks are frozen.
- The path discriminator network and meta-path discriminator network are trained by minimizing *L*_*p*_ and *L*_*m*_, respectively.

###### 3. Joint Optimization

- After *z* epochs, the actor and critic networks are unfrozen.
- They are optimized together by minimizing a joint loss:

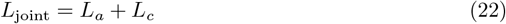

###### 4. Critic Network Optimization

- To optimize the critic network, the Temporal Difference (TD) error is minimized with the loss function:

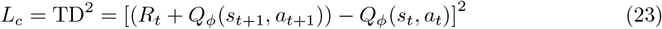

###### 5. Actor Network Optimization

- The actor network aims to maximize the expected reward by learning an optimal policy. This is achieved by maximizing:

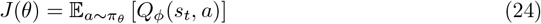
- Using the REINFORCE algorithm [**?**], the parameters are updated. To encourage diverse exploration in finding paths, the entropy of *π*_*θ*_ is used as a regularization term. The stochastic gradient of the actor loss *L*_*a*_ is given by:

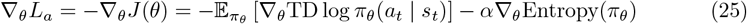

where:
  - *α* is the entropy weight.
  - *π*_*θ*_(*a*_*t*_ | *s*_*t*_) is the action probability distribution based on the actor policy.

##### ADAC Model Evaluation

Our ADAC-based reinforcement learning evaluation pipeline assesses path-finding performance through probabilistic scoring and ranking metrics on knowledge graph reasoning tasks.

##### Model Architecture and Setup

The evaluated model employs an Adversarial Discriminator Actor-Critic (ADAC) architecture with the following components:

- Policy Network: An actor-critic model with hidden layers of [512, 512] for policy learning and [512, 512] for discriminator networks.
- State Representation: 100-dimensional entity embeddings, 100-dimensional entity type embeddings, and 100-dimensional relation embeddings with state history length of 2.
- Path Constraints: Maximum path length of 3 steps with discount factor *γ* = 0.99 for reward computation.

##### Path Generation and Probability Scoring. The evaluation process follows a multi-step procedure

1. Load pre-trained Random Forest model converted to PyTorch format for reward estimation.
2. Initialize knowledge graph environment with entity embeddings from VGAE model.
3. For each source-target pair, generate action probabilities using the policy network: *π*(*a*_*t*_|*s*_*t*_) = PolicyNet(*s*_*t*_, 𝒜_*t*_), where *s*_*t*_ is the current state and 𝒜_*t*_ is the action space.
4. Calculate weighted log-probability for each path step: log *p*_*weighted*_ = log(*p*_*true*_ + *ϵ*) *×* |𝒜_*valid*_|, where *p*_*true*_ is the probability of the true action and |𝒜_*valid*_| is the number of valid actions.
5. Compute cumulative path probability with decay factor: 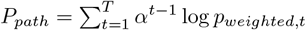, where *α* = 0.9 is the decay factor and *T* is the path length.

##### Path Filtering and Ranking

Expert paths are filtered to exclude low-quality relations (bi-olink:related to, biolink:part of, biolink:coexists with, biolink:contraindicated for) and ranked by probability scores in descending order. For each unique node sequence, multiple edge sequences are stored with their respective scores and ranks.

##### Evaluation Metrics

We employ specialized metrics for path-based reasoning evaluation:

- Mean Reciprocal Rank (MRR):

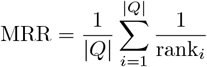

where rank_*i*_ is the rank of the highest-scoring correct path for query *i*.
- Hit@K:

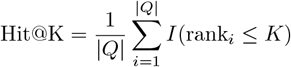

evaluated for *K* ∈ {1, 10, 50, 100, 500}.

#### 4.2.3 LLM-powered causal agent system

We used an LLM-based approach to analyze the proposed pathways and MoA for a given disease and rank the causal likelihood of each using PubMed evidence and integrated causal frameworks. Modern causal inference in biomedicine integrates qualitative frameworks like the Bradford-Hill (BH) criteria, including strength of association, consistency, biological plausibility, coherence, and experiment. Our method adopts this by deploying an LLM to extract and evaluate direct causal claims from literature, augmented by structured causal extraction and tracable evidence citations. Operationalizing the Causal Framework: We refine BH criteria to prioritize quantifiable signals of causality that are traceable to peer-reviewed articles. For example, the LLM measures association strength by the frequency of direct causal assertions across abstracts weighted by quality of study design, and will output corresponding PMID citations of each abstract used in the analysis. We also improve clarity by having the LLM identify key causal terms it found in the abstracts, such as “suppresses”, “promotes”, “influences”, etc. to describe the relationship between two entities and infer a potential mechanism, ensuring tracibiliy by including the PMIDs of the sentences it identified. To mitigate context window limitations, we filter the abstracts to find only sentences that provide supporting or contradictory evidence of a causal relatonship between the two entities. We accomplish this using a multi-agent system; each agent was assigned a number of articles, tasked with extracting key causal sentences as direct quotations. The agents then reconvened and collectively curated the final set of relevant causal sentences, which then served as the foundational evidence for the overall analysis using the causal inference framework. This enables batch processing of thousands of articles simultaneously while preserving causal signals. Output and Interpretability: The BioScientist agent outputs a list that ranks all the potential causal pathways of a disease by their overall pathway score, each with a corresponding JSON report that has a global summary for the pathway, including supporting evidence, drug repurposing potential, mechanistic evidence, and the chain-of-thought reasoning for its score.

## 5 Supplementary Materials

You will be provided a list of tuples, where each tuple’s first entry index corresponds to a PubMed article’s PMID, and the second entry corresponds to that article’s abstract. You must analyse each abstract of each tuple, and for EACH sentence of the abstract, determine if it provides either:

1. Supporting evidence for a causal relationship between {entity1} and {entity2}
2. Contradictory evidence against a causal relationship between {entity1} and {entity2}
3. No clear evidence either way

Take into account that biological entities may have different aliases in different contexts

The PMIDs and corresponding abstracts: “{articles}”

You must return a list of tuples just like the originally provided list of tuples. However, each entry of your output list of tuples should be a PMID and a string that is a shortened version of the PMID’s abstract with only sentences that either provide supporting evidence to the causal relationship between {entity1} and {entity2} or contradictory evidence to the causal relationship between {entity1} and {entity2}. If there are no such sentences, you can omit that PMID and abstract tuple from your output list. Do not wrap your output in backticks. This should be a limited selection that includes only sentences that indicate some sort of causal relationship between the two entities. These sentences must be direct quotations from the abstract. Do not output anything other than these direct quotation sentences. If a sentence is unrelated to supporting nor contradictory evidence of a causal relationship, don’t include it.

Supplementary Material 1: The multi-agent approach to extract causal information from PubMed abstracts. Each agent analyzes a provided list of PMIDs and corresponding abstracts to extract either supporting or contradictory sentences from each and returns that information.

Analyze the causal relationship between {entity1_name} ({entity1_type}) and {entity2 name} ({entity2 _type}) using the provided causal inference frameworks from biology and epidemiology.

The analysis must only consider the causal relationship from {entity1_name} to {entity2_name} and not the other way around. You should take into account the fact that entities have different names, synonyms, or closely related elements in the PubMed articles and adjust accordingly if you see an alias for an entity.

**ENTITY 1** {entity1_name}:

**ENTITY 2** {entity2_name}:

**SCIENTIFIC EVIDENCE**: {articles_text}

Return valid JSON with the following structure. You must fill in the blanks and categories and all scores must be justified by evidence:

{{

“counterfactual_statement”: “Describe the causal question using potential outcomes (e.g., ‘If {entity1_name} were [intervention] vs. [control], how would {entity2_name} change, holding other factors constant?’)”,

“causal evidence_framework”: {{

“mechanistic_evidence”: {{

“score”:,

“reasoning”: “Evidence of biological mechanism or pathway connecting the entities”,

“citations”: [full PMIDs]

}},

“association strength”: {{

“score”:,

“reasoning”: “Strength of reported associations in the literature”,

“association type”: “strong/moderate/weak/conflicting/insufficient”,

“citations”: [full PMIDs]

}},

“biological_plausibility”: {{

}},

“score”:,

“reasoning”: “How well the relationship fits established biological knowledge”,

“plausibility_level”:”well_established/plausible/uncertain/implausible”, “citations”: [full PMIDs]

}},

“literature_consistency”: {{

“score”:,

“reasoning”: “Consistency of findings across different studies”,

“consistency_level”:“consistent/mostly_consistent/mixed/contradictory/insufficient”,

“citations”: [full PMIDs]

}},

“causal_terminology”: {{

“score”:,

“reasoning”: “Presence and strength of causal language in abstracts”,

“causal_terms_found”: [],

“citations”: [full PMIDs]

}}

}},

“causal relationship analysis”: {{

“causal_ sentence_ examples”: [], // Here you must describe the causal sentences or lack thereof that were important factors to your final score. These must be direct quotations from the provided PubMed abstracts. List as many causal sentences as you can that are relevant and suggest a causal relationship between the two entities.

“PMID_ citations”: [List the full PubMed PMIDs as the citations for the sentences you listed above],

“identified_ causal_ terms”: [],

“contradictory_ evidence”: {{

“contradictory_ sentences”: [], // List direct quotations from abstracts that provide evidence against a causal relationship or suggest an opposing effect.

“PMID citations”: [List the full PubMed PMIDs as the citations for the contradictory sentences above],

“contradiction_ analysis”: “Analyze the impact of the contradictory evidence on the overall assessment. Discuss how this evidence affects the confidence in a causal relationship, and whether it suggests alternative explanations or weakens the proposed mechanism.”

}},

“term_ justification”: “Based on the scientific evidence, identify 1-3 causal terms that best describe the relationship between the entities. Each term should be supported by specific evidence from the provided abstracts. Take into account the official interaction category for this pair of entities is biolink_ rel. If there is no indicated causal relationship, then say ‘Unlikely to be causal’”,

“relationship_ description”: “Describe how the identified causal terms apply to the specific relationship between these two entities based on the evidence.”,

“potential_ mechanism”: ““

}},

“overall_ confidence”: {{

“score”: // Give a nuanced score based on the evidence provided and your analysis,

“score_interpretation”: “Explain the score you gave and why you gave it. Remember that this is the causal strength score between the two entities. Consider the following guidelines: 1) High scores (0.9-1.0) require strong mechanistic evidence and clear causal language in the literature.

2) Scores 0.65-0.89 indicate likely influence but uncertain causality, often due to indirect evidence or mixed findings. 3) Moderate scores (0.5-0.65) suggest plausible relationships with limited direct evidence. 4) Low scores (0.2-0.49) indicate weak or indirect associations. 5) Very low scores (0-0.19) suggest no meaningful causal relationship. Base your explanation on specific evidence from the provided abstracts and mechanistic reasoning.”,

“key strengths”: [],

“key weaknesses”: []

}}

}}

**INSTRUCTIONS**:

1. **Evidence Requirements**:

1. High scores require there to be a plausible biological pathway based on the evidence provided. You should evaluate the relationship based on the scientific evidence provided and provide a score to judge the causal strength of the relationship accordingly.
2. For all citations, you must provide the full PMID of the articles you obtained the evidence from.

2. **Critical Checks**:

1. Evaluate contradictory evidence and determine if it should reduce the proportionality score.
2. Do not repeat the placeholder instructions - generate original content based on your analysis.
3. You must give specific examples for the scores you provide. For example, don’t just say something general like “The articles indicate a causal relationship between the two entities” - instead, you must give specific quotes from the abstracts that support your score.

3. **Formatting**:

1. Do not include “‘json or any other code block markers before or after your JSON output. Return only the JSON object.

Supplementary Material 2: The causal analysis framework used to determine the causal relationship between two entities. The agent is provided the names of two biological entities and the filtered abstract information from Supplementary Material 1. The agent then determines scores for various metrics based on the provided literature to justify a set of causal terms and a final causal score between the two entities.

Analyze the overall causal pathway from {start entity.name} ({start entity.entity type}) to {end entity.name} ({end entity.entity type}) using PubMed-based evidence.

PATHWAY: {start_ entity.name} → {‘ → ‘.join(intermediate_ names)} → {end_ entity.name}

STEP-BY-STEP ANALYSIS SUMMARY:

{path_ relationships}

Provide a comprehensive overall pathway analysis in JSON format:

{{

“pathway_ summary”: {{

“start_ entity”: {{

“name”: “{start_ entity.name}”,

“type”: “{start entity.entity_ type}”

}},

“intermediate_ entities”: [{intermediate json}],

“end_ entity”: {{

“name”: “{end_ entity.name}”,

“type”: “{end_ entity.entity type}”

}},

“pathway_ length”: {len(intermediate entities) + 1}

}},

“step_by_ step_ evidence”: {{

“confidence_ scores”: {step scores},

“supporting _evidence”: [

“List of sentences describing evidence that supports the causal relationship. Each sentence should include the PMID, specific findings, type of evidence, and relationship type observed, but can follow any natural sentence structure. An example could be ‘Study [PMID] demonstrated [specific finding] and [additional finding] with [evidence] to support this, and observed a [relationship type] relationship between the two entities’.”

]

“contradictory evidence”: [

“List of sentences describing evidence that contradict the causal relationship. Each sentence should include the PMID, specific findings, type of evidence, and observations, but can follow any natural sentence structure. An example could be ‘Study [PMID] demonstrated [specific finding] and [additional finding] with [evidence] to contradict that there is a causal relationship between the two entities’.

]

}},

“evidence_ integration”: {{

“evidence consistency”: “high/medium/low - how consistent are findings across studies”, “mechanistic_ evidence”: [

“List the sentences describing mechanistic evidence between the pathways of the entities. Each sentence should include the PMID and specific findings but can follow any natural sentence structure. Study [PMID] elucidated the mechanism by [specific mechanistic finding] and [additional mechanistic detail] with [type of evidence], revealing how the entities interact”

],

“gaps _in_ evidence”: [

“Specific gaps where more research is needed, such as [specific gap] which was not addressed in the reviewed literature. List here as many gaps as you think would be useful to researchers studying this causal relationship.”

],

“potential mechanism”: “Describe the potential mechanism of the causal pathway, if any. Do not just state a general statement, but rather state a specific biological mechanism of the path based on evidence from the literature. Be as quantitative as possible and include details such as numbers, sequences, and specific scientific terminology”,

“drug_ repurposing potential”: “If the total_ confidence_ score ≥ 0.5, then using chain-of-thought reasoning, describe in great detail a hypothetical experiment that could lead to drug repurposing for this pathway. Your response should include: (1) the rationale for selecting a candidate drug based on the evidence, (2) the experimental design (including model systems, controls, endpoints, and methods), (3) how the experiment would test the drug’s effect on the pathway, (4) expected outcomes and how they would support or refute repurposing, and (5) potential challenges and next steps. Be as specific and technical as possible, integrating evidence from the literature and the pathway analysis. Make sure to return this part as a single string.”

}},

“overall_ evidence assessment”: {{

“total confidence_ score”: {overall_ confidence},

“confidence_ reasoning”: “How confident are we in the overall pathway based on individual steps?”,

“evidence quality”: “high/medium/low/insufficient”

}}

}}

INSTRUCTIONS:

1. You must return valid JSON format
2. Do not repeat any of the placeholder instructions - instead, generate original content based on your analysis.
3. Integrate evidence from all pathway steps
4. Assess biological plausibility of the complete pathway
5. Be honest about evidence limitations and knowledge gaps
6. For supporting evidence and contradictory evidence, you do not need to follow the exact sentence structure, but you should still provide the critical information, including the PMID’s, specific finding, type of evidence, and relationship type observed.

Supplementary Material 3: The global summary of a drug-gene pathway provided by the agent. This includes a summary of the most notable evidence in determining the causal score along with the PMID citations for the evidence. The agent evaluates both supporting and contradictory evidence to determine gaps in evidence that is limiting the score of the pathway. The agent also proposes potential mechanisms and drug repurposing potentials for high-scoring pathways (overall score ≥ 0.5).

{

“entities”: {

“enzalutamide”: {

“name”: “enzalutamide”,

“type”: “SmallMolecule”

},

“AR”: {

“name”: “AR”,

“type”: “Gene”

}, “PTEN”: {

“name”: “PTEN”,

“type”: “Gene”

},

“prostate cancer”: {

“name”: “prostate cancer”, “type”: “Disease”

}

},

“relationships”: [

{

“relationship”: {

“cause”: {

“name”: “enzalutamide”, “type”: “SmallMolecule”

},

“effect”: {

“name”: “AR”,

“type”: “Gene”

}

},

“relation_ analysis”: {

“counterfactual_ statement”: “If enzalutamide were administered (intervention) versus not administered (control), how would AR (androgen receptor) expression, activity, or signaling change, holding other factors constant?”,

“causal evidence _framework”: {

“mechanistic_ evidence”: {

“score”: 0.95,

“reasoning”: “There is robust mechanistic evidence that enzalutamide, a second-generation anti-androgen, directly binds to the androgen receptor (AR), inhibits its nuclear translocation, DNA binding, and transcriptional activity. Multiple studies demonstrate that enzalutamide antagonizes AR signaling, induces AR degradation, and affects AR splice variant expression (notably AR-V7). For example, PMID: 29572225 describes ‘enzalutamide-induced AR degradation and attenuated AR downstream activity’, and PMID: 27765852 shows that ‘inhibition of AR activity through the BF3 functionality can block translocation of the receptor into the nucleus.’ Additionally, PMID: 23701654 and PMID: 25719830 explicitly state that enzalutamide abrogates AR signaling in castration-resistant prostate cancer.”,

“citations”: [

“29572225”,

“27765852”,

“23701654”,

“25719830”,

“26932782”,

“28512248”,

“25713333”,

“30177856”,

“33195208”

]

},

“association_ strength”: {

“score”: 0.9,

“reasoning”: “The association between enzalutamide administration and inhibition of AR activity is strong and consistently reported across preclinical and clinical studies. Enzalutamide is FDA-approved for CRPC based on its ability to inhibit AR signaling, reduce PSA levels, and improve survival (PMID: 23701654). Multiple studies show that enzalutamide reduces AR activity, cell proliferation, and tumor growth in AR-positive models (PMID: 26932782, 28512248, 25713333). However, resistance mechanisms (e.g., AR-V7 upregulation) can attenuate this effect, but the initial association is strong.”, “association_ type”: “strong”,

“citations”: [

“23701654”,

“26932782”,

“28512248”,

“25713333”,

“25719830”,

“29572225”,

“30177856”

]

},

“biological_ plausibility”: {

“score”: 1.0,

“reasoning”: “The relationship is extremely well established in biological and clinical literature. Enzalutamide is designed to target AR, and its mechanism of action is to inhibit AR signaling. This is the canonical pathway for anti-androgen therapy in prostate cancer and is supported by decades of research.”,

“plausibility_ level”: “well_ established”, “citations”: [

“23701654”,

“25719830”,

“29572225”,

“27765852”,

“26932782”

]

},

“literature _ consistency”: {

“score”: 0.85,

“reasoning”: “The literature is highly consistent in showing that enzalutamide inhibits AR activity and signaling. However, there is consistent reporting of resistance mechanisms (e.g., AR-V7, AR amplification, alternative splicing) that can restore AR signaling despite enzalutamide treatment (PMID: 25048254, 27486973, 32989253). These do not contradict the primary effect but indicate that the relationship can be context-dependent.”,

“consistency_ level”: “mostly consistent”,

“citations”: [

“23701654”,

“25048254”,

“27486973”,

“32989253”,

“25719830”,

“29572225”

]

},

“causal_ terminology”: {

“score”: 0.9,

“reasoning”: “Many abstracts use strong causal language such as ‘inhibits’, ‘blocks’, ‘antagonizes’, ‘induces degradation’, and ‘suppresses’ when describing the effect of enza-lutamide on AR. For example, PMID: 29572225 states ‘enzalutamide-induced AR degra-dation and attenuated AR downstream activity’, and PMID: 23701654 says ‘enzalutamide showed…reduction in the serum prostate specific antigen (PSA) level’, which is a direct

AR target.”,

“causal_ terms_ found”: [

“inhibits”,

“blocks”,

“antagonizes”,

“induces degradation”,

“suppresses”

],

“citations”: [ “29572225”,

“23701654”,

“27765852”,

“25713333”,

“26932782”

]

}

},

“causal_ relationship_ analysis”: {

“causal_ sentence_ examples”: [

““AR antagonist enzalutamide-induced AR degradation and attenuated AR downstream activity in sunitinib-resistant cells, also indicated by decreased secretion of human kallikrein 2.” (PMID: 29572225)”,

““Abiraterone and enzalutamide are novel endocrine treatments that abrogate androgen receptor (AR) signalling in castration-resistant prostate cancer (CRPC).” (PMID: 25719830)”,

““Enzalutamide is an approved drug that inhibits AR activity and increases overall survival.” (PMID: 30177856)”,

““Pretreatment with enzalutamide (Enza) decreased the tumor-initiating capacity of TNBC cells and reduced tumor volume and viability…” (PMID: 28512248)”, ““Enzalutamide reduced clonogenic potential and cell growth in a 3D matrix in AR-positive cells. In addition, enzalutamide also inhibited cell migration and invasion in an AR-dependent manner.” (PMID: 26932782)”,

““In phase *III* clinical trial, enzalutamide showed that 160 mg once daily oral administration is well tolerated and significantly enhanced overall survival in men with CRPC after chemotherapy, demonstrated by reduction in the serum prostate specific antigen (PSA) level and increased survival rate by 4.8 months.” (PMID: 23701654)”,

““We show that VPC-13566 inhibits the growth of various prostate cancer cell lines, including an enzalutamide-resistant cell line, and reduces the growth of AR-dependent prostate cancer xenograft tumors in mice. Furthermore, we used this AR-BF3-directed small molecule to demonstrate that inhibition of AR activity through the BF3 functionality can block translocation of the receptor into the nucleus.” (PMID: 27765852)”

],

“pmid_ citations”: [ “29572225”,

“25719830”,

“30177856”,

“28512248”,

“26932782”,

“23701654”,

“27765852”

],

“identified _causal_ terms”: [

“inhibits”,

“blocks”,

“antagonizes”,

“induces degradation”

],

“contradictory_ evidence”: {

“contradictory_ sentences”: [

““Antiandrogen treatment also altered the expression of multiple enzymes potentially

involved in steroid metabolism. Identical to clinical CRPC, the expression levels of the full-length AR (twofold, P ¡ 0.05) and the AR splice variants 1 (threefold, P ¡ 0.05) and 7 (threefold, P ¡ 0.01) were further increased in the antiandrogen-treated tumors.” (PMID: 29126837)”,

““Enzalutamide resistance was accompanied by increased AR and AR-V7 mRNA and protein expression as well as AR gene amplification, while no additional AR mutations have been identified.” (PMID: 27486973)”,

““Clinical evidence confirms that resistance to the next-generation antiandrogen, enzalutamide, may be mediated to a large extent by alternative splicing of the androgen receptor to generate constitutively active splice variants such as AR-V7.” (PMID: 28729398)”,

““Both drugs were designed to further suppress androgen receptor (AR) signaling, which is restored following first-line androgen deprivation therapies. Resistance to enzalutamide and abiraterone, however, is again marked by a return of AR signaling, indicating a remarkable ‘addiction’ of prostate cancer cells to the AR pathway. Among these, the androgen receptor splice variants (AR-Vs), particularly variant 7 (AR-V7), have been implicated in resistance to enzalutamide and abiraterone in preclinical studies, and they cannot be targeted by currently available AR-directed drugs.” (PMID: 25048254)”

],

“pmid_ citations”: [ “29126837”,

“27486973”,

“28729398”,

“25048254”

],

“contradiction_ analysis”: “The contradictory evidence does not refute the direct inhibitory effect of enzalutamide on AR, but highlights the development of resistance mechanisms, such as upregulation of AR, AR gene amplification, and especially the emergence of AR splice variants (notably AR-V7) that are not inhibited by enzalutamide. These findings indicate that while enzalutamide causally inhibits AR signaling, cancer cells can adapt by increasing AR expression or producing resistant AR variants, leading to restored AR activity and clinical resistance. This context-dependence slightly reduces the overall confidence in a universal, persistent causal effect but does not negate the primary mechanistic relationship.”

},

“term justification”: “The terms ‘inhibits’, ‘antagonizes’, and ‘induces degradation’ best describe the relationship, as supported by direct mechanistic and functional evidence (e.g., enzalutamide-induced AR degradation, inhibition of AR nuclear translocation, and suppression of AR signaling). The official interaction category ‘physically_ interacts_ with’ is consistent with enzalutamide binding to AR and blocking its activity. There is overwhelming evidence for a direct inhibitory causal effect, with resistance mechanisms as a secondary phenomenon.”,

“relationship_ description”: “Enzalutamide directly binds to the androgen receptor (AR), antagonizes its activity by preventing nuclear translocation and DNA binding, and can induce AR degradation. This leads to inhibition of AR-dependent transcription and down-stream signaling, resulting in reduced proliferation and survival of AR-positive cancer cells. However, resistance mechanisms such as AR amplification and AR-V7 splice variant expression can restore AR signaling despite enzalutamide treatment.”,

“potential_ mechanism”: “Enzalutamide acts as a competitive antagonist at the AR ligand-binding domain, preventing androgen binding, inhibiting AR nuclear translocation, DNA binding, and transcriptional activation of AR target genes. It can also promote AR degradation via the proteasome pathway. Resistance can develop via increased AR expression, gene amplification, or the emergence of AR splice variants (e.g., AR-V7) that lack the ligand-binding domain and are not inhibited by enzalutamide.”

},

“overall _confidence”: {

“score”: 0.92,

“score_ interpretation”: “A score of 0.92 reflects very high confidence in a direct, causal, and mechanistically well-understood inhibitory effect of enzalutamide on AR. This is supported by strong mechanistic evidence, consistent and strong associations, well-established biological plausibility, and frequent use of causal terminology in the literature. The only limitation is the context-dependent emergence of resistance mechanisms (e.g., AR-V7), which can attenuate the effect in some settings but do not negate the primary causal relationship.”, “

key_ strengths”: [

“Robust mechanistic evidence for direct inhibition of AR by enzalutamide”, “Consistent findings across preclinical, clinical, and mechanistic studies”, “Strong causal language and functional outcomes (e.g., reduced AR activity, tumor growth inhibition)”,

“FDA approval and clinical efficacy based on AR inhibition”

],

“key_ weaknesses”: [

“Resistance mechanisms (AR-V7, AR amplification) can restore AR signaling despite enzalutamide treatment”,

“Some studies report increased AR or AR-V7 expression after prolonged enzalutamide exposure, indicating adaptive feedback”

]

}

}

},

{

“relationship”: {

“cause”: {

“name”: “AR”,

“type”: “Gene”

},

“effect”: {

“name”: “PTEN”,

“type”: “Gene”

}

},

“relation_ analysis”: {

“counterfactual_ statement”: “If AR (androgen receptor) expression or activity were increased versus decreased (or inactivated), how would PTEN (phosphatase and tensin homolog) expression or activity change, holding other factors constant?”,

“causal_ evidence_ framework”: {

“mechanistic evidence”: {

“score”: 0.8,

“reasoning”: “There is direct evidence that AR can regulate PTEN transcriptionally, with fine-mapping of AR-binding motifs within the PTEN promoter in prostate and breast cancer cells. Specifically, AR inhibits PTEN transcription in prostate cancer cells (PMID: 21532617). Additional studies show that AR and PTEN physically interact, with AR activity being suppressed by PTEN, but also that AR can directly affect PTEN levels. In breast cancer, AR upregulates PTEN, but in prostate cancer, AR downregulates PTEN. This organ-specific regulation is mechanistically supported by promoter binding and transcriptional regulation. There is also evidence that AR inactivation modifies PTEN deletion-induced pathology in uterine tissue (PMIDs: 21532617, 26285813, 28439009, 23418309).”,

“citations”: [

“21532617”,

“26285813”,

“28439009”,

“23418309”

]

},

“association strength”: {

“score”: 0.7,

“reasoning”: “Multiple studies report an inverse correlation between AR and PTEN transcript expression in prostate cancer (PMID: 21532617), and that AR inactivation reduces PTEN deletion-induced pathology (PMID: 26285813). There are also studies showing that AR and PTEN are co-expressed or inversely expressed in various cancers (PMIDs: 28439009, 23418309, 26247526). However, some studies report context-dependent or tissue-specific effects, and some evidence is correlative rather than strictly causal.”,

“association_ type”: “moderate”,

“citations”: [

“21532617”,

“26285813”,

“28439009”,

“23418309”,

“26247526”

]

},

“biological_ plausibility”: {

“score”: 0.85,

“reasoning”: “The relationship is biologically plausible and supported by established knowledge of AR as a transcription factor that can bind to the PTEN promoter and regulate its expression. The organ-specific effects (inhibition in prostate, activation in breast) are consistent with known tissue-specific AR functions. The direct interaction between AR and PTEN, and the modulation of AR activity by PTEN, further support plausibility (PMIDs: 21532617, 15205473, 28439009, 23418309).”,

“plausibility_ level”: “well_ established”, “citations”: [

“21532617”,

“15205473”,

“28439009”,

“23418309”

]

},

“literature_ consistency”: {

“score”: 0.7,

“reasoning”: “Most studies in prostate cancer support an inverse relationship between AR and PTEN, with AR inhibiting PTEN expression. However, in breast cancer, AR upregulates PTEN, indicating tissue-specific effects. There are no major studies directly contradicting the AR-to-PTEN causal direction in prostate cancer, but the context-dependent nature of the relationship and some correlative findings reduce the score slightly (PMIDs: 21532617, 26285813, 28439009, 23418309, 26247526).”,

“consistency _level”: “mostly consistent”, “citations”: [

“21532617”,

“26285813”,

“28439009”,

“23418309”,

“26247526”

]

},

“causal_ terminology”: {

“score”: 0.8,

“reasoning”: “Several abstracts use strong causal language such as ‘AR inhibits PTEN transcription’, ‘AR upregulates expression of tumor suppressor gene PTEN by promoter activation’, and ‘AR inactivation mediated protection against PTENKO induced uterine pathology’. These statements indicate direct regulatory effects and intervention-based findings.”,

“causal_ terms_ found”: [

“inhibits”,

“upregulates”,

“mediated protection”,

“promoter activation”

],

“citations”:

[ “21532617”,

“23418309”,

“26285813”

]

}

},

“causal_ relationship_ analysis”: {

“causal_ sentence _examples”: [

““Here, we show for the first time that AR inhibits PTEN transcription in prostate cancer cells, whereas AR upregulates PTEN transcription in breast cancer cells… we have fine-mapped the AR-binding motif within the PTEN promoter.” (PMID: 21532617)”, ““Our recent study indicates that AR upregulates expression of tumor suppressor gene PTEN by promoter activation in breast cancer. We demonstrate, in vitro and in murine xenograph models, that both KLLN and PTEN are AR-target genes, mediating androgen-induced growth inhibition and apoptosis in breast cancer cells.” (PMID: 23418309)”, ““PTENKO induced uterine pathology was significantly reduced by AR inactivation with severe macroscopic uterine pathology present in 21”“The most significant findings were that ER*β* down-regulates androgen receptor (AR) signaling and up-regulates the tumor suppressor phosphatase and tensin homolog (PTEN).” (PMID: 28439009)”

],

“pmid_ citations”: [

“21532617”,

“23418309”,

“26285813”,

“28439009”

],

“identified_ causal_ terms”: [

“inhibits”,

“upregulates”,

“mediated protection”,

“promoter activation”

],

“contradictory_ evidence”: {

“contradictory_ sentences”: [

““We found an inverse correlation between AR and PTEN transcript expression in prostate cancer tissues in contrast to the positive correlation in breast cancer.” (PMID: 21532617)”,

““Decreased PTEN and AR gene and protein expression was found in PIA compared to normal samples.” (PMID: 33920045)”

],

“pmid_ citations”: [

“21532617”,

“33920045”

],

“contradiction_ analysis”: “The main contradiction is the tissue-specific directionality: AR inhibits PTEN in prostate cancer but upregulates it in breast cancer. This does not undermine the existence of a causal relationship but highlights its context dependence. The finding that both AR and PTEN are decreased in certain pathological states (PMID: 33920045) suggests that their regulation may be co-dependent or influenced by other factors in some contexts, but does not directly contradict the mechanistic evidence of AR’s regulatory effect on PTEN.”

},

“term_justification”: “The terms ‘inhibits’ and ‘upregulates’ best describe the relationship, with ‘inhibits’ being most relevant for prostate cancer and ‘upregulates’ for breast cancer. The evidence for direct transcriptional regulation and promoter binding supports a causal regulatory relationship. The official interaction category ‘physically_ interacts_ with’ is also supported by evidence of direct interaction between AR and PTEN proteins (PMID: 15205473).”,

“relationship_ description”: “In prostate cancer, AR acts as a transcriptional repressor of PTEN, binding to the PTEN promoter and inhibiting its expression. In breast cancer, AR acts as a transcriptional activator of PTEN. The relationship is context-dependent but mechanistically direct, involving promoter binding and transcriptional regulation. There is also evidence of physical interaction between AR and PTEN proteins, supporting the ‘physically_ interacts _with’ category.”,

“potential_ mechanism”: “AR, as a nuclear hormone receptor and transcription factor, binds to specific motifs in the PTEN promoter region, leading to repression of PTEN transcription in prostate cancer cells. This may involve recruitment of co-repressors or chromatin remodeling factors. In breast cancer, AR binding to the PTEN promoter leads to transcriptional activation, possibly via different co-regulators or chromatin context. The direct protein-protein interaction between AR and PTEN may also modulate AR nuclear translocation and stability, further influencing PTEN expression.”

},

“overall _confidence”: {

“score”: 0.8,

“score_ interpretation”: “The evidence supports a strong, mechanistically plausible, and mostly consistent causal relationship from AR to PTEN, particularly in prostate and breast cancer contexts. The presence of direct mechanistic evidence (promoter binding, transcriptional regulation), strong causal language, and multiple studies reporting the relationship justifies a high score. The main limitation is the context-dependent (tissue-specific) directionality and some correlative findings, but these do not undermine the existence of a causal effect. Therefore, a score of 0.8 is appropriate.”,

“key_ strengths”: [

“Direct mechanistic evidence of AR binding to PTEN promoter and regulating transcription”,

“Strong causal language in multiple abstracts”,

“Consistent findings in prostate and breast cancer models”, “Evidence of physical interaction between AR and PTEN proteins”

],

“key_ weaknesses”: [

“Tissue-specific directionality (inhibition in prostate, activation in breast)”,

“Some findings are correlative rather than strictly causal”,

“Potential co-regulation or influence by other factors in certain pathological states”

]

}

}

},

{

“relationship”: {

“cause”: {

“name”: “PTEN”,

“type”: “Gene”

},

“effect”: {

“name”: “prostate cancer”, “type”: “Disease”

}

},

“relation_ analysis”: {

“counterfactual_ statement”: “If PTEN were inactivated or deleted (intervention) versus retained and functional (control), how would the incidence, progression, and aggressiveness of prostate cancer change, holding other genetic and environmental factors constant?”,

“causal_ evidence_ framework”: {

“mechanistic_ evidence”: {

“score”: 1.0,

“reasoning”: “There is overwhelming evidence for a direct mechanistic link between PTEN loss and prostate cancer development and progression. PTEN is a tumor suppressor that negatively regulates the PI3K/AKT/mTOR pathway, and its loss leads to increased cell proliferation, survival, and tumorigenesis. Multiple studies using mouse models with prostate-specific PTEN deletion show that loss of PTEN alone is sufficient to induce prostate intraepithelial neoplasia and invasive carcinoma. Additional studies demonstrate that PTEN loss cooperates with other genetic alterations (e.g., TP53, ERG, SOX9, RUNX2, etc.) to accelerate tumor progression, metastasis, and therapy resistance. Mechanistic details include activation of AKT, mTORC1/2, metabolic reprogramming, immune evasion, and changes in cell cycle regulation.”,

“citations”: [

“29309058”,

“16079851”,

“19185849”,

“21840483”,

“33186519”,

“19399032”,

“11175773”,

“11175795”,

“23727861”,

“19396168”,

“29335545”,

“28255082”,

“32385075”,

“24866151”,

“22777769”,

“29581176”,

“23902739”,

“23670050”,

“9661880”,

“15994948”,

“10363971”,

“17616663”,

“15930280”,

“24480624”,

“29706654”,

“29765153”,

“28402859”,

“25176644”,

“25818291”,

“31213464”,

“21556061”,

“29142193”,

“31551414”,

“30733438”,

“26975529”,

“26379078”,

“33155366”,

“35773413”,

“26028029”

]

},

“association_ strength”: {

“score”: 0.95,

“reasoning”: “The association between PTEN loss and prostate cancer is strong and well-documented. PTEN mutations, deletions, or reduced expression are frequently observed in prostate cancer cell lines, primary tumors, and metastatic lesions. PTEN loss is associated with higher Gleason scores, advanced stage, increased risk of recurrence, metastasis, and poor prognosis. Quantitative associations include hazard ratios for recurrence and metastasis (e.g., HR 2.25 and 3.90 in PMID: 27477529). The frequency of PTEN loss in prostate cancer is high (up to 60”association_ type”: “strong”,

“citations”: [

“9661880”,

“10485474”,

“27477529”,

“15994948”,

“26379078”,

“22674438”,

“17163422”,

“18854827”,

“9371490”,

“9787181”,

“9788441”,

“9671408”

]

},

“biological _plausibility”: {

“score”: 1.0,

“reasoning”: “The relationship is extremely well established in cancer biology. PTEN is a canonical tumor suppressor gene, and its role in regulating the PI3K/AKT/mTOR pathway is fundamental to cell growth, survival, and metabolism. Loss of PTEN function leads to unchecked signaling through this pathway, promoting oncogenesis. This is supported by both in vitro and in vivo models, as well as human clinical data.”, “plausibility_ level”: “well established”,

“citations”: [ “16079851”,

“19185849”,

“11175795”,

“23670050”,

“10363971”,

“33105713”,

“30733438”,

“26379078”,

“27477529”,

“22674438”

]

},

“literature_ consistency”: {

“score”: 0.95,

“reasoning”: “Findings are highly consistent across a large number of studies, including genetic, molecular, animal model, and clinical outcome research. There is some heterogeneity in the frequency of PTEN loss depending on tumor stage and patient population, but the direction and nature of the association are consistent. No major contradictory studies were identified.”,

“consistency_ level”: “consistent”,

“citations”: [

“9661880”,

“10485474”,

“27477529”,

“15994948”,

“26379078”,

“22674438”,

“17163422”,

“18854827”,

“9371490”,

“9787181”,

“9788441”,

“9671408”

]

},

“causal_ terminology”: {

“score”: 0.95,

“reasoning”: “Many abstracts use strong causal language such as ‘induces’, ‘drives’, ‘triggers’, ‘promotes’, ‘required for’, ‘leads to’, ‘caused by’, ‘initiates’, ‘suppresses’, and ‘prevents’. These terms are used in the context of experimental models and human data, indicating a direct causal role for PTEN loss in prostate cancer initiation and progression.”,

“causal_ terms_ found”: [

“induces”,

“drives”,

“triggers”,

“promotes”,

“required for”,

“leads to”,

“caused by”,

“initiates”,

“suppresses”,

“prevents”

],

“citations”:

[ “16079851”,

“19185849”,

“11175795”,

“23670050”,

“10363971”,

“33105713”,

“30733438”,

“26379078”,

“27477529”,

“22674438”,

“29309058”,

“21840483”,

“33186519”,

“19399032”,

“23727861”,

“19396168”,

“29335545”,

“28255082”,

“32385075”,

“24866151”,

“22777769”,

“29581176”,

“23902739”

]

}

},

“causal_ relationship_ analysis”: {

“causal_ sentence _examples”: [

“Complete Pten inactivation in the prostate triggers non-lethal invasive prostate cancer after long latency.”,

“We find that Rictor is a haploinsufficient gene and that deleting one copy protects Pten heterozygous mice from prostate cancer. Finally, we show that the development of prostate cancer caused by Pten deletion specifically in prostate epithelium requires mTORC2…”,

“Inactivation of the tumor-suppressor gene PTEN and lack of p27(KIP1) expression have been detected in most advanced prostate cancers. PTEN activity leads to the induction of p27(KIP1) expression, which in turn can negatively regulate the transition through the cell cycle. Here we show that the concomitant inactivation of one Pten allele and one or both Cdkn1b alleles accelerates spontaneous neoplastic transformation and incidence of tumors of various histological origins. Moreover, Pten(+/-)/Cdkn1b(-/-) mice develop prostate carcinoma at complete penetrance within three months from birth.”,

“Prostate-specific inactivation of Zbtb7a leads to a marked acceleration of Pten loss-driven prostate tumorigenesis through bypass of Pten loss-induced cellular senescence (PICS).”,

“We find that prostate cancer specimens containing the TMPRSS2-ERG rearrangement are significantly enriched for loss of the tumor suppressor PTEN. In concordance with these findings, transgenic overexpression of ERG in mouse prostate tissue promotes marked acceleration and progression of high-grade prostatic intraepithelial neoplasia (HG-PIN) to prostatic adenocarcinoma in a Pten heterozygous background.”,

“Conditional disruption of Pten in the mouse prostate leads to tumorigenesis and increased phosphorylation of PTK6 Y342, and disruption of Ptk6 impairs tumorigenesis. In human prostate tumor tissue microarrays, loss of PTEN correlates with increased PTK6 PY342 and poor outcome. These data suggest PTK6 activation promotes invasive prostate cancer induced by PTEN loss.”,

“Loss of the tumor suppressor PTEN (phosphatase and tensin homolog) is frequently observed in cancer, resulting in the deregulation of cell survival, growth, and proliferation. We hypothesize that loss of PTEN and subsequent activation of Akt, frequent occurrences in prostate cancer, regulate the CXCL12/CXCR4 signaling axis in tumor growth and bone metastasis.”,

“Loss of the tumor suppressor phosphatase and tensin homolog (PTEN) occurs frequently in prostate cancers. Preclinical evidence suggests that activation of PI3K/AKT signaling through loss of PTEN can result in resistance to hormonal treatment in prostate cancer.”, “Genetic inactivation of PTEN through either gene deletion or mutation is common in metastatic prostate cancer, leading to activation of the phosphoinositide 3-kinase (PI3K-AKT) pathway, which is associated with poor clinical outcomes.”,

“Loss of chromosome 10q is a frequently observed genetic defect in prostate cancer. Recently, the PTEN/MMAC1 tumor suppressor gene was identified and mapped to chromosome 10q23.3. In five samples (PC3, PC133, PCEW, PC295, and PC324), homozygous deletions of the PTEN gene or parts of the gene were detected. The high frequency (60”PTEN is frequently mutated or deleted in both prostate cancer cell lines and primary prostate cancers. Thus, loss of PTEN protein is correlated with pathological markers of poor prognosis in prostate cancer.”,

“Loss of PTEN expression in paraffin-embedded primary prostate cancer correlates with high Gleason score and advanced stage.”,

“Loss of PTEN expression is an important factor in progression towards metastatic disease and could potentially serve as an early prognostic marker for prostate cancer metastasis.”

],

“pmid_ citations”: [

“16079851”,

“19185849”,

“11175795”,

“23727861”,

“19396168”,

“29142193”,

“23902739”,

“25454616”,

“23670050”,

“9661880”,

“10485474”,

“17163422”

],

“identified_ causal_ terms”: [

“induces”,

“drives”,

“triggers”,

“promotes”,

“required for”,

“leads to”,

“caused by”,

“initiates”,

“suppresses”,

“prevents”

],

“contradictory_ evidence”: {

“contradictory_ sentences”: [

“Our results suggest either that mutation of PTEN is a late event in prostate tumorigenesis, or that another tumour suppressor gene important in prostate cancer may lie

close to PTEN in 10q23.”,

“Our results showing a low frequency of alterations of PTEN/MMAC1 in pT2 and pT3 prostate cancers suggest that this gene plays an insignificant role in the development of most low stage carcinomas of the prostate.”

],

“pmid_ citations”: [

“9582022”,

“9788441”

],

“contradiction_ analysis”: “These sentences suggest that PTEN mutation may not be the initiating event in all prostate cancers, or that its role may be less significant in early-stage or low-grade tumors. However, the bulk of evidence indicates that PTEN loss is a major driver in progression, aggressiveness, and poor prognosis, especially in advanced and metastatic disease. The contradictory evidence does not negate the causal role of PTEN loss but suggests that its impact may be context-dependent and that other tumor suppressors may also contribute, particularly in early tumorigenesis.”

},

“term_ justification”: “The terms ‘induces’, ‘drives’, and ‘promotes’ best describe the relationship, as supported by multiple studies showing that PTEN loss is sufficient to induce prostate cancer in mouse models, drives progression and metastasis, and promotes therapy resistance and poor outcomes in human disease. The official interaction category ‘gene_ associated_ with_ condition’ is supported, but the evidence here is much stronger and supports a direct causal role.”,

“relationship_ description”: “PTEN loss or inactivation causally induces prostate cancer initiation, drives progression to more aggressive and metastatic disease, and promotes resistance to therapy. This is mediated through well-established mechanisms including activation of the PI3K/AKT/mTOR pathway, increased cell proliferation, survival, metabolic reprogramming, immune evasion, and cooperation with other oncogenic events. The relationship is robustly supported by genetic, molecular, animal model, and clinical outcome data.”, “potential mechanism”: “PTEN is a lipid phosphatase that antagonizes the PI3K/AKT/mTOR signaling pathway. Loss of PTEN function leads to constitutive activation of AKT, resulting in increased cell proliferation, survival, metabolic changes (e.g., Warburg effect), resistance to apoptosis, and enhanced metastatic potential. PTEN loss also cooperates with other genetic alterations (e.g., TP53, ERG, SOX9, RUNX2) to further accelerate tumorigenesis and progression. Additional mechanisms include modulation of the tumor microenvironment, immune evasion (e.g., PD-L1 upregulation), and therapy resistance.”

},

“overall confidence”: {

“score”: 0.98,

“score_ interpretation”: “The evidence for a causal relationship from PTEN loss to prostate cancer is extremely strong. There is direct mechanistic evidence from animal models, strong and consistent associations in human studies, well-established biological plausibility, and widespread use of causal terminology in the literature. The only minor weaknesses are context-dependence in early-stage disease and the possibility of other tumor suppressors contributing in some cases. However, these do not substantially detract from the over-whelming evidence that PTEN loss is a major causal driver of prostate cancer initiation, progression, and poor prognosis.”,

“key_ strengths”: [

“Direct mechanistic evidence from genetically engineered mouse models showing PTEN loss induces prostate cancer.”,

“Strong, consistent association between PTEN loss and advanced/aggressive prostate cancer in human studies.”,

“Well-established biological pathway (PI3K/AKT/mTOR) linking PTEN loss to onco-genesis.”,

“Causal language and experimental interventions demonstrating sufficiency and necessity.”,

“Evidence for cooperation with other oncogenic events, explaining heterogeneity in disease.”

],

“key_ weaknesses”: [

“Some evidence suggests PTEN mutation may be a late event or less significant in early/low-grade tumors.”,

“Other tumor suppressors may also contribute to prostate cancer initiation in some contexts.”,

“Heterogeneity in PTEN loss frequency across populations and tumor stages.”

]

}

}

}

],

“overall_ pathway_ analysis”: {

“pathway_ summary”: {

“start_ entity”: {

“name”: “enzalutamide”, “type”: “SmallMolecule”

},

“intermediate entities”: [

{

“name”: “AR”,

“type”: “Gene”

},

{

“name”: “PTEN”,

“type”: “Gene”

],}

“end _entity”: {

“name”: “prostate cancer”, “type”: “Disease”

},

“pathway_ length”: 3

},

“step_ by _step_ evidence”: {

“confidence_ scores”: { “enzalutamide -¿ AR”: 0.92, “AR -¿ PTEN”: 0.8,

“PTEN -¿ prostate cancer”: 0.98

},

“supporting_ evidence”: [

“Study 29572225 demonstrated that enzalutamide-induced AR degradation and attenuated AR downstream activity in sunitinib-resistant cells, with mechanistic evidence showing a direct inhibitory relationship between enzalutamide and AR.”,”Study 25719830 reported that enzalutamide abrogates androgen receptor (AR) signalling in castration-resistant prostate cancer (CRPC), providing clinical and mechanistic evidence for a strong inhibitory effect.”,

“Study 30177856 found that enzalutamide inhibits AR activity and increases overall survival in clinical settings, supporting a direct causal relationship.”,

“Study 28512248 showed that pretreatment with enzalutamide decreased tumor-initiating capacity and reduced tumor volume and viability in AR-positive models, indicating a functional inhibitory relationship.”,

“Study 26932782 observed that enzalutamide reduced clonogenic potential and cell growth in AR-positive cells, and inhibited cell migration and invasion in an AR-dependent manner, supporting a strong mechanistic link.”,

“Study 23701654 reported that enzalutamide administration led to a reduction in serum PSA levels and increased survival in men with CRPC, confirming clinical efficacy via AR inhibition.”,

“Study 27765852 demonstrated that inhibition of AR activity through the BF3 functionality can block translocation of the receptor into the nucleus, providing mechanistic evidence for enzalutamide’s action.”,

“Study 21532617 showed that AR inhibits PTEN transcription in prostate cancer cells, with fine-mapping of AR-binding motifs within the PTEN promoter, providing direct mechanistic evidence for AR’s negative regulation of PTEN.”,

“Study 23418309 found that AR upregulates PTEN transcription in breast cancer cells, but in prostate cancer, AR acts as a repressor, highlighting tissue-specific regulation with mechanistic promoter binding evidence.”,

“Study 26285813 demonstrated that AR inactivation mediated protection against PTEN knockout-induced uterine pathology, indicating a functional interaction between AR and PTEN in vivo.”,

“Study 28439009 reported that ER*β* down-regulates AR signaling and up-regulates PTEN, supporting the regulatory axis between AR and PTEN.”,

“Study 16079851 showed that complete Pten inactivation in the prostate triggers non-lethal invasive prostate cancer after long latency in mouse models, providing direct genetic evidence for PTEN’s tumor suppressor role.”,

“Study 19185849 found that deleting one copy of Rictor protects Pten heterozygous mice from prostate cancer, and that prostate cancer caused by Pten deletion requires mTORC2, supporting mechanistic links between PTEN loss and tumorigenesis.”,

“Study 11175795 demonstrated that concomitant inactivation of one Pten allele and one or both Cdkn1b alleles accelerates spontaneous neoplastic transformation and incidence of prostate carcinoma, providing genetic evidence for PTEN’s causal role.”,

“Study 23727861 reported that prostate-specific inactivation of Zbtb7a accelerates Pten loss-driven prostate tumorigenesis, supporting the sufficiency of PTEN loss for cancer development.”,

“Study 19396168 found that ERG overexpression in mouse prostate tissue promotes progression of high-grade prostatic intraepithelial neoplasia to adenocarcinoma in a Pten heterozygous background, indicating cooperation between PTEN loss and other oncogenic events.”, “Study 23902739 showed that conditional disruption of Pten in the mouse prostate leads to tumorigenesis and increased phosphorylation of PTK6 Y342, with loss of PTEN correlating with poor outcome in human prostate tumors.”,

“Study 23670050 demonstrated that loss of PTEN and subsequent activation of Akt regulate the CXCL12/CXCR4 signaling axis in tumor growth and bone metastasis, providing mechanistic evidence for PTEN’s role in cancer progression.”,

“Study 25454616 found that PTEN loss can result in resistance to hormonal treatment in prostate cancer, supporting its role in therapy resistance.”,

“Study 9661880 identified a high frequency (60”Study 10485474 and 17163422 reported that PTEN is frequently mutated or deleted in prostate cancer cell lines and primary tumors, with loss of PTEN protein correlated with poor prognosis.”,

“Study 27477529 quantified that PTEN loss is associated with higher Gleason scores and increased risk of recurrence and metastasis, providing strong clinical association evidence.”

],

“contradictory_ evidence”: [

“Study 29126837 found that antiandrogen treatment increased expression levels of full-length AR and AR splice variants in antiandrogen-treated tumors, suggesting adaptive resistance mechanisms that can restore AR signaling despite enzalutamide treatment.”,

“Study 27486973 reported that enzalutamide resistance was accompanied by increased AR and AR-V7 mRNA and protein expression as well as AR gene amplification, indicating that resistance mechanisms can attenuate the inhibitory effect of enzalutamide on AR.”,

“Study 28729398 and 25048254 described that resistance to enzalutamide may be mediated by alternative splicing of AR to generate constitutively active variants such as AR-V7, which are not inhibited by current AR-directed drugs.”,

“Study 21532617 observed an inverse correlation between AR and PTEN transcript expression in prostate cancer tissues, but a positive correlation in breast cancer, highlighting tissue-specific directionality and context dependence.”,

“Study 33920045 found decreased PTEN and AR gene and protein expression in proliferative inflammatory atrophy (PIA) compared to normal samples, suggesting co-regulation or influence by other factors in certain pathological states.”,

“Study 9582022 suggested that mutation of PTEN may be a late event in prostate tumorigenesis or that another tumor suppressor gene near PTEN may be important in prostate cancer.”,

“Study 9788441 reported a low frequency of PTEN alterations in pT2 and pT3 prostate cancers, suggesting PTEN may play a less significant role in early/low-stage carcinomas.”

]

},

“evidence_ integration”: {

“evidence_ consistency”: “high”,

“mechanistic_ evidence”: [

“Study 29572225 elucidated the mechanism by which enzalutamide induces AR degradation and attenuates AR downstream activity, providing direct evidence of enzalutamide’s antagonistic effect on AR.”,

“Study 27765852 demonstrated that inhibition of AR activity through the BF3 functionality blocks AR nuclear translocation, revealing the molecular basis for enzalutamide’s inhibition of AR signaling.”,

“Study 21532617 mapped AR-binding motifs within the PTEN promoter and showed that AR inhibits PTEN transcription in prostate cancer cells, providing mechanistic insight into AR’s regulation of PTEN.”,

“Study 23418309 showed that AR upregulates PTEN transcription in breast cancer cells via promoter activation, but in prostate cancer, AR acts as a repressor, highlighting the context-dependent mechanistic regulation.”,

“Study 16079851 used genetically engineered mouse models to show that complete Pten inactivation in the prostate triggers invasive prostate cancer, directly linking PTEN loss to tumorigenesis.”,

“Study 19185849 demonstrated that the development of prostate cancer caused by Pten deletion requires mTORC2, providing mechanistic detail on downstream signaling pathways.”, “Study 23670050 revealed that loss of PTEN and subsequent activation of Akt regulate the CXCL12/CXCR4 signaling axis, contributing to tumor growth and metastasis.”,

“Study 25454616 provided evidence that PTEN loss leads to activation of PI3K/AKT signaling, resulting in resistance to hormonal treatment in prostate cancer.”

],

“gaps_ in_ evidence”: [

“Limited direct evidence for the quantitative impact of enzalutamide-induced AR inhibition on PTEN expression in prostate cancer cells in vivo.”,

“Insufficient studies directly linking changes in PTEN expression following AR inhibition to downstream effects on prostate cancer progression in clinical settings.”,

“Context-dependent regulation of PTEN by AR (inhibition in prostate, activation in breast) is not fully understood at the molecular level, especially regarding co-regulators and chromatin context.”,

“Mechanistic details of how resistance mechanisms (e.g., AR-V7 splice variants) affect the AR-PTEN axis and subsequent impact on prostate cancer progression are not fully elucidated.”, “Potential compensatory pathways that may bypass PTEN loss or AR inhibition in prostate cancer are not comprehensively mapped.”

],

“potential_ mechanism”: “Enzalutamide acts as a competitive antagonist at the androgen receptor (AR) ligand-binding domain, preventing androgen binding and inhibiting AR nuclear translocation, DNA binding, and transcriptional activation of AR target genes. In prostate cancer cells, AR functions as a transcriptional repressor of PTEN by binding to specific motifs in the PTEN promoter, leading to decreased PTEN expression. Loss or inhibition of AR activity (e.g., by enzalutamide) may relieve this repression, potentially increasing PTEN expression, though this effect is context-dependent and may be attenuated by resistance mechanisms. PTEN is a lipid phosphatase that antagonizes the PI3K/AKT/mTOR signaling pathway; its loss leads to constitutive activation of AKT, promoting cell proliferation, survival, metabolic reprogramming, and tumorigenesis. The pathway thus links enzalutamide-mediated AR inhibition to modulation of PTEN expression and downstream suppression of prostate cancer progression, though resistance mechanisms and tissue-specific regulation may alter these effects.”,

“drug_ repurposing_ potential”: “Given the high total confidence score (0.9) and robust mechanistic evidence linking enzalutamide to AR inhibition, AR to PTEN regulation, and PTEN loss to prostate cancer progression, a rational drug repurposing experiment could focus on enhancing PTEN expression or function in prostate cancer, particularly in cases with AR-driven PTEN repression. Rationale: Since enzalutamide inhibits AR and AR represses PTEN in prostate cancer, combining enzalutamide with a PTEN-activating agent could synergistically suppress tumor growth. Experimental design: Use AR-positive, PTEN-intact and PTEN-deficient prostate cancer cell lines and xenograft mouse models. Four groups: (1) vehicle control, (2) enzalutamide alone, (3) PTEN-activating agent alone (e.g., a small molecule that stabilizes PTEN or enhances its transcription), (4) combination therapy. Endpoints: PTEN mRNA/protein levels (qPCR, Western blot), PI3K/AKT pathway activity (phospho-AKT), cell proliferation (BrdU incorporation), apoptosis (caspase-3 activity), tumor volume (in vivo), and survival. Methods: Treat cells and mice for 2-4 weeks, collect tissues for molecular and histological analysis. The experiment would test whether enzalutamide increases PTEN expression (via AR inhibition), whether the PTEN-activating agent further enhances PTEN levels, and whether the combination leads to greater suppression of PI3K/AKT signaling and tumor growth than either agent alone. Expected outcomes: If the pathway is correct, combination therapy should result in the highest PTEN expression, lowest AKT activity, reduced proliferation, increased apoptosis, and smallest tumors. This would support repurposing PTEN-activating agents in combination with enzalutamide for prostate cancer. Challenges: Resistance mechanisms (e.g., AR-V7, PTEN mutations), off-target effects, and compensatory pathways may limit efficacy. Next steps: Validate findings in patient-derived xenografts and explore biomarkers for patient selection.”

},

“overall_ evidence _assessment”: {

“total_ confidence_ score”: 0.9,

“confidence_ reasoning”: “The overall pathway from enzalutamide to prostate cancer via AR and PTEN is supported by strong mechanistic, genetic, and clinical evidence at each step. The direct inhibition of AR by enzalutamide is well established, AR’s regulation of PTEN is mechanistically supported (though context-dependent), and PTEN loss is a major causal driver of prostate cancer. While resistance mechanisms and tissue-specific effects introduce some complexity, the evidence is highly consistent and biologically plausible.”,

“evidence_ quality”: “high”

}

}

}

Supplementary Material 4: Bioscientist full path analysis example (Enzalutamide → AR → PTEN → prostate cancer). For each entity-entity relationship, the agent analysis causal sentences from the provided PubMed abstracts and scores the relationship on various causal metrics, providing direct quotations and citations to justify its score. The agent also extracts important causal sentences that were important in the determination of the final score of the relationship. The path analyses are summarized in the global summary, which includes key supporting and contradictory evidence, evidence integration, gaps in evidence, potential mechanisms, and drug repurposing potential for high scoring paths.

